# Scavenging dicarbonyls with 5’-O-pentyl-pyridoxamine increases HDL net cholesterol efflux capacity and attenuates atherosclerosis and insulin resistance

**DOI:** 10.1101/529339

**Authors:** Jiansheng Huang, Huan Tao, Patricia G. Yancey, Zoe Leuthner, Linda S. May-Zhang, Ju-Yang Jung, Youmin Zhang, Lei Ding, Venkataraman Amarnath, Dianxin Liu, Sheila Collins, Sean S. Davies, MacRae F. Linton

## Abstract

**Objective:** Oxidative stress contributes to the development of insulin resistance (IR) and atherosclerosis. Peroxidation of lipids produces reactive dicarbonyls such as Isolevuglandins (IsoLG) and malondialdehyde (MDA) that covalently bind plasma/cellular proteins, phospholipids, and DNA leading to altered function and toxicity. We examined whether scavenging reactive dicarbonyls with 5’-O-pentyl-pyridoxamine (PPM) protects against the development of IR and atherosclerosis in *Ldlr*^-/-^ mice.

**Methods:** Male or female *Ldlr*^-/-^ mice were fed a western diet (WD) for 16 weeks and treated with PPM versus vehicle alone. Plaque extent, dicarbonyl-lysyl adducts, efferocytosis, apoptosis, macrophage inflammation, and necrotic area were measured. Plasma MDA-LDL adducts and the in vivo and in vitro effects of PPM on the ability of HDL to reduce macrophage cholesterol were measured. Blood Ly6C^hi^ monocytes and ex vivo 5-ethynyl-2’-deoxyuridine (EdU) incorporation into bone marrow CD11b+ monocytes and CD34+ hematopoietic stem and progenitor cells (HSPC) were also examined. IR was examined by measuring fasting glucose/insulin levels and tolerance to insulin/glucose challenge.

**Results:** PPM reduced the proximal aortic atherosclerosis by 48% and by 46% in female and male *Ldlr*^-/-^ mice, respectively. PPM also decreased IR and hepatic fat and inflammation in male *Ldlr*^-/-^ mice. Importantly, PPM decreased plasma MDA-LDL adducts and prevented the accumulation of plaque MDA- and IsoLG-lysyl adducts in *Ldlr*^-/-^ mice. In addition, PPM increased the net cholesterol efflux capacity of HDL from *Ldlr*^-/-^ mice and prevented both the in vitro impairment of HDL net cholesterol efflux capacity and apoAI crosslinking by MPO generated hypochlorous acid. Moreover, PPM decreased features of plaque instability including decreased proinflammatory M1-like macrophages, IL-1β expression, myeloperoxidase, apoptosis, and necrotic core. In contrast, PPM increased M2-like macrophages, Tregs, fibrous cap thickness, and efferocytosis. Furthermore, PPM reduced inflammatory monocytosis as evidenced by decreased blood Ly6C^hi^ monocytes and proliferation of bone marrow monocytes and HSPC from *Ldlr*^-/-^ mice.

**Conclusions:** PPM has pleotropic atheroprotective effects in a murine model of familial hypercholesterolemia, supporting the therapeutic potential of reactive dicarbonyl scavenging in the treatment of IR and atherosclerotic cardiovascular disease.

## 1. INTRODUCTION

Atherosclerosis is a systemic inflammatory disorder that initiates early in life and develops in the arterial wall over decades. Clinical and mechanistic studies have shown that many factors facilitate the development of atherosclerosis, presenting potential targets for therapeutic modification. Although clinical trials of lowering LDL-C have consistently been demonstrated to reduce cardiovascular events, a considerable residual risk of cardiovascular disease still exists, which may be due in part to inflammation [1–3]. This residual inflammatory risk highlights the need to find novel therapeutic approaches for targeting atherosclerosis to achieve more effective reductions in cardiovascular risk.

Pleotropic functions of HDL, including anti-inflammatory function and mediation of reverse cholesterol transport (RCT), are central in protecting against cardiovascular disease [4]. Cholesterol efflux capacity, a measure of HDL function in the first step of RCT, was reported to be inversely associated with measures of cardiovascular events, independent of HDL-C levels [5–7]. Recent observations support the concept that qualitative changes in HDL composition promote its atheroprotective functions [8, 9]. HDL is also identified as an important target for treatment of type 2 diabetes (T2D), which is associated with increased risk for atherosclerotic cardiovascular events. HDL promotes the uptake of glucose in skeletal muscle and increases the synthesis and secretion of insulin from pancreatic β cells and thus may have beneficial effects on glycemic control [10, 11]. Clinical studies found that reconstituted HDL increased plasma insulin and decreased plasma glucose in patients with T2D by activating AMP-activated protein kinase [11, 12]. Thus, mounting evidence demonstrates that HDL function is crucial in chronic inflammatory disorders, such as T2D and atherosclerosis.

Studies have shown that myeloperoxidase (MPO) promotes the formation of dysfunctional HDL and there is abundant dysfunctional HDL in human atherosclerotic lesions [13–15]. Several reactive dicarbonyls including malondialdehyde (MDA), 4-oxo-nonenal (ONE), and isolevuglandins (IsoLG) are endogenously formed during lipid peroxidation. Immunization with MDA epitopes leads to atheroprotection in mouse and rabbit models [16, 17], highlighting the importance of MDA epitopes in atherosclerosis. Synthetic MDA-lysine can promote monocyte activation and vascular complications via induction of inflammatory pathways [18]. Modifying apoAI or HDL by MDA or IsoLG blocks cholesterol efflux from macrophages [19–21]. In addition to lipid dicarbonyl modification, myeloperoxidase (MPO) mediated chlorination of tyrosine and oxidation of tryptophan in apoAI impairs cholesterol efflux [14, 15]. Furthermore, dysfunctional apoAI is present in human atherosclerotic lesions including MDA modified HDL and chlorinated and oxidized apoAI residues [14, 15, 21]. Interestingly, our recent studies showed that IsoLG, MDA, and ONE modification of HDL reduces its anti-inflammatory function in macrophages treated with lipopolysaccharide (LPS) [19, 20, 22]. Importantly, HDL from subjects with familial hypercholesterolemia (FH) versus normocholesterolemic controls has reduced cholesterol efflux capacity and anti-inflammatory function and contains significantly more IsoLG, MDA, and ONE adducts, which likely contribute to the dysfunction of FH-HDL [19, 20, 22]. In addition, HDL from humans with coronary artery disease versus control subjects have increased MDA adducts, which results in reduced endothelial anti-inflammatory and repairing effects [23]. Taken together, mounting evidence supports the concept that HDL atheroprotective function can be impaired and that reactive dicarbonyls such as MDA, ONE and IsoLG and other oxidative modifications of apoAI contribute to HDL dysfunction, insulin resistance (IR), and the development of atherosclerosis.

In the current studies, we utilized a potent reactive dicarbonyl scavenger, 5’-O-pentyl-pyridoxamine (PPM), to dissect the role of reactive carbonyl species in causing HDL dysfunction in promoting the development of IR and atherosclerosis. Our earlier studies demonstrated that, compared to lysine residues, PPM reacts nearly 2000 times faster with the reactive dicarbonyl, IsoLG, thereby preventing lysine modification [24]. We found that scavenging reactive dicarbonyls with PPM prevents MPO-mediated impairment of HDL cholesterol efflux capacity in vitro, and, importantly, in vivo administration of PPM to hypercholesterolemic mice increased HDL cholesterol efflux capacity without affecting plasma cholesterol levels. Consistent with the effects on HDL function, administration of PPM to *Ldlr*^-/-^ mice consuming a western diet reduced IR, atherosclerotic lesion extent, lesion pro-inflammatory macrophages, blood Ly6C^hi^ monocytosis, and lesion efferocytosis. Furthermore, PPM treatment increased the plaque fibrous cap thickness and reduced the necrotic area, thereby promoting features of plaque stability. Taken together, our results suggest that reactive dicarbonyl scavenging has high therapeutic potential in improving inflammation, preventing HDL dysfunction, IR, and atherosclerosis development.

## 2. MATERIAL AND METHODS

### 2.1. Mice

*Apoe^-/-^ and Ldlr*^-/-^ mice were purchased from Jackson Laboratories (Bar Harbor, ME). Mice were maintained in micro-isolator cages with *ad libitum* access to a rodent chow diet containing 4.55 % fat (PMI 5010, St. Louis, Mo) and water. The Institutional Animal Care and Use Committee of Vanderbilt University Medical Center approved mouse protocols. Experimental procedures and animal care were performed according to the regulations of Vanderbilt University’s Institutional Animal Care and Usage Committee.

### 2.2. Treatment of *Ldlr*^-/-^ mice with PPM to examine the impact on atherosclerosis

5’-O-pentyl-pyridoxamine (PPM) was solubilized in water and prepared as a 100 mM stock and stored as small aliquots at −80°C until use. Fresh working solutions were prepared before each assay and diluted in water to the appropriate concentrations. As recommended by the scientific statement on animal atherosclerosis studies from the American Heart Association, studying the atherosclerosis in both male and female mice is a standard procedure to evaluate potential gender specific contributions to atherosclerosis. Therefore, for the atherosclerosis studies, female and male *Ldlr*^-/-^ mice on a western diet (WD) were treated with water alone or with 1 g/L of PPM in the drinking water for 16 weeks after pretreatment of mice on chow diet with PPM or 2 weeks. The % by weight composition of the WD (Harlan, TD.88137) used in the experiments was 19.5% Casein, 17.3 % protein, 0.3% DL-methionine, 34% sucrose, 21% anhydrous milk fat, 0.2% cholesterol, 5% cellulose (fiber), 3.5% mineral mix (AIN-76, 170915), 0.4% calcium carbonate, 1% vitamin mix (Teklad,40060), 15% corn starch, and 0.004% ethoxyquin.

### 2.3. Reagents

Fresh human HDL was isolated from normal subjects by sequential density ultracentrifugation as previously described [20]. Maloncarbonyl bis-(dimethylacetal) (cat#: 820756) and H_2_O_2_ (cat#: 216763) were purchased from Sigma-Aldrich (St. Louis, MO). The chemiluminescent Western Lightning ultra-reagent was obtained from PerkinElmer Inc. (Waltham, MA). Primary antibody to apoAI (cat#: NB600-609) CCR2 (cat#: NBP2-67700) were purchased from Novus Biologicals. Primary anti-MOMA2 (cat#: MCA519G) was purchased from BioRad. Secondary antibody HRP conjugated-rabbit anti-goat IgG (cat#: AD106P) and primary anti-MPO antibody (cat#: AF3667) were purchased from Sigma and R&D systems, respectively. Arg1 antibody (cat#: GTX109242) was purchased from GeneTex. Secondary fluorescent antibodies (FITC goat-anti-rabbit Secondary antibody, cat#: A11034; AF568 Donkey-anti-rabbit Secondary antibody, cat#: A10042; AF488 Donkey-anti-rat Secondary antibody, cat#: A21208) for immunofluorescence staining were obtained from Invitrogen.

### 2.4. Effect of PPM on modification of HDL with MDA

HDL modification with MDA was done in 50 mM phosphate buffer (pH 7.0) containing 100 μM DTPA. For the PPM inhibition reaction, 1 mM PPM or 5 mM PPM was incubated with 1mg/mL HDL (cat#: J64903) for 10 min and then 250 μM MDA was added to initiate the reaction. The reactions were carried out at 37 °C for 24 h and stopped by dialysis against PBS overnight.

### 2.5. Effect of PPM on modification of HDL with MPO

The MPO-mediated HDL modification reactions were done in phosphate buffer (50 mM, pH 7.0) containing 100 μM DTPA, 1 mg/ml of HDL protein (cat#: J64903), 70 nM purified human MPO (Athens Research, cat#: 16-14-130000, A430/A280 ratio of 0.79), and 100 mM NaCl [25]. For the PPM inhibition reaction, 1 mM PPM was incubated with 1mg/mL HDL for 10 min and then 70 nM MPO, 160 μM H_2_O_2_, and 100 mM NaCl were added to initiate the reaction. The reactions were incubated at 37 °C for 1 h. In all reactions, working solutions of H_2_O_2_ were made fresh daily by diluting 30% H_2_O_2_ according to the extinction coefficient for H_2_O_2_ at 240 nm, 39.4 M^−1^cm^−1^ [26].

### 2.6. Effect of PPM on crosslinking of Lipid-Free ApoAI with Hypochlorous Acid

The hypochlorous acid (HOCl)-mediated ApoAI modification reactions were performed in 50 mM phosphate buffer (pH 7.0) containing 100 μM diethylenetriaminepentaacetic acid. For the PPM inhibition reaction, PPM was incubated with 0.2 mg/mL ApoAI for 10 min and then 50 μM HOCl were added to initiate the reaction. The reactions were carried out at 37°C for 1 h. The samples were immediately used for western blot to determine the effect of PPM on crosslinking of lipid-free ApoAI.

### 2.7. Isolation of peritoneal macrophages

Peritoneal macrophages were isolated 4 days after intraperitoneal injection of 5% of thioglycolate. Peritoneal cells were collected by rinsing with Ca^2+^ and Mg^2+^ free PBS and then centrifuged and resuspended in DMEM containing 10% heat-inactivated FBS (Sigma Aldrich, cat#: B207151) and 100 units/ml penicillin/streptomycin. Cells were plated onto 24-well culture plates (1*10^6^ cells/well) and allowed to adhere for 2 h. Nonadherent cells were removed by washing two times with DPBS, and adherent macrophages were used for experiments.

### 2.8. Serum lipid measurements, FPLC lipoprotein profile, and ApoB-depleted serum preparation

Serum total cholesterol and triglyceride (TG) levels were measured by the enzyme-based method (Fisher, cat#: 890339 RC) after overnight fasting. Lipid profiling of FPLC-separated lipoprotein fractions was done using gel filtration analysis with a Superose 6 column on a Waters 600 FPLC system. Mouse serum (100μl) was injected onto the column and separated with buffer containing 0.15 M NaCl, 0.01 M Na_2_HPO4, and 0.1mM EDTA (pH 7.5), at a flow rate of 0.5 mL/min. Cholesterol content of the fractions was determined using an enzymatic assay (Fisher, cat#: 891069 RA) [27]. ApoB-depleted serum was prepared by adding 200 μL of PEG 8,000 (20% in 200 mM glycine) to 500 μL serum [28, 29]. After incubation for 15 min at room temperature, and then centrifugation at 1,900g for 20 min, the supernatant containing the apoB-depleted serum was collected and used for cholesterol efflux experiments [30].

### 2.9. Measurement of HDL capacity to reduce macrophage cholesterol

To measure the capacity of HDL to reduce macrophage cholesterol content, an enzymatic cholesterol assay was used as described [31, 32]. Briefly, *Apoe*^-/-^ macrophages were incubated for 48 h with DMEM containing acetylated LDL (Alfa aesar, cat#: J65029, 40 μg/ml). The cells were then washed and incubated for 1 h in DMEM containing 0.1% BSA. The cells were then washed and incubated for 24 h in DMEM supplemented with HEPES in the presence of 50 μg/mL human HDL, modified HDL, or 2.5% apoB-depleted serum from mice. Cellular cholesterol was measured after incubation with DMEM alone or with cholesterol acceptor using an enzymatic cholesterol assay [31, 32]. Briefly, isopropanol/NP-40 (vol/vol:9/1) was added to the dried isopropanol lipid extracts and vortexed for 10 sec. Then 40 ul of each sample was added to 96-well plates and catalase solutions (Sigma, cat#:C1345) in 50 mM potassium phosphate buffer were added to the wells for both standards and samples to lower the background. Then, 100 ul of the enzyme and fluorescent probe mixture (ADHP, cat#: AnaSpec # 85500; cholesterol oxidase, Cat#: C8649-100U; cholesterol esterase, Cat#:C9281-100UN; horseradish peroxidase, Cat#: P8375-1kun) were prepared in reagent A buffer (sodium chloride, cholic acid, Triton-100 in 100 mM potassium phosphate buffer) and added into each well. The fluorescence intensity (excitation wavelength 530nm, emission wavelength 580nm) was measured immediately after incubation at 37°C for 25mins. The cholesterol efflux capacity was calculated as ((cellular cholesterol with DMEM alone - cellular cholesterol with acceptor)/cellular cholesterol with DMEM alone * 100).

### 2.10. Western blotting

Samples were incubated with β-mercaptoethanol and loading buffer for 10 min at 55 °C and then resolved by NuPAGE Bis-Tris electrophoresis using 4-12% gels for 1.5 h. The proteins were transferred onto PVDF membranes (Amersham Bioscience) at 150 V for 1.5 h. Membranes were blocked with 5% milk at room temperature for 2 h and incubated with primary antibodies specific to human apoAI or MDA overnight at 4°C. The secondary antibodies conjugated with HRP were incubated with the membranes for 2 h at room temperature. Protein bands were visualized with ECL western blotting detection reagents (GE Healthcare) and quantified using ImageJ software (NIH).

### 2.11. Glucose tolerance test (GTT) and insulin tolerance test (ITT)

Insulin levels were measured by ELISA (Mercodia, cat#: 10-1113-01). For the GTT, mice were fasted for 4h, and 10 μL blood was taken from the tail vein for measurement of baseline levels. Mice were then injected i.p. with 1g/kg of glucose. Blood samples were then taken at 15, 30, 45, 60, 90, 120 minutes and analyzed for blood glucose with a clinical meter [33]. For ITT, mice were fasted for 4h, and after taking a baseline blood sample, the mice were injected intraperitoneally with insulin (0.5U/kg). Blood samples were then taken at 15, 30, 60, 90, 120 minutes and analyzed for blood glucose with a clinical meter [33]. The homeostatic model assessment for insulin resistance (HOMA-IR) was calculated using the equation fasting insulin (μU/ml) × fasting plasma glucose (mmol/liter)/22.5.

### 2.12. Hematoxylin and eosin (H&E) staining of liver tissue and NAFLD severity score

H&E staining of liver tissue was performed as previously described [34]. Briefly, livers were immediately embedded in paraffin, sectioned, and mounted on glass microscope slides. The sections were stained with hematoxylin and eosin and examined using light microscopy. Nonalcoholic fatty liver disease severity score, which estimates the severity of fatty liver by the summation of steatosis, lobular inflammation, and hepatocyte balloon degeneration, was evaluated semi-quantitatively: based on the severity of steatosis, lobular inflammation and hepatocellular ballooning.

### 2.13. Measurement of atherosclerotic lesions and immunofluorescence analyses

We followed the guidelines for development and execution of experimental design and interpretation of atherosclerosis in animal studies [35]. The mean lesion area from 15 serial sections was used for the aortic root atherosclerotic lesion size per mouse as described [36]. Immunofluorescence staining was performed using sections that were 40–60□μm distal of the aortic sinus. Briefly, aortas were flushed with PBS through the left ventricle, and the entire aorta was dissected and en face was stained for the neutral lipid using Sudan IV as described [36]. The extent of atherosclerosis was examined by Oil-Red-O staining of neutral lipids in crosssections of the proximal aorta (15 alternate 10 μm cryosections for each mouse aorta). Images of the proximal and distal aorta lesions were analyzed using the KS300 imaging system (Kontron Electronik GmbH) [37, 38]. MDA was detected by immunofluorescence staining using anti-MDA primary antibody (FITC, Abcam#: ab27615). Immunofluorescence staining was used to examine the levels of MOMA2, MPO, arginase 1 (Arg1), and C-C chemokine receptor 2 (CCR2) in atherosclerotic lesions [37]. For each mouse, two sections were stained, and quantitation was performed on the entire cross section of all stained sections. Briefly, 5 μm cross-sections of the proximal aorta were fixed in cold acetone (Sigma), blocked with 5% goat serum dilution (GSD), and incubated overnight at 4°C with primary antibody to MOMA2, MPO, Arg1, or CCR2. After incubation for 1h at 37 °C with the fluorescent labeled secondary antibody, the nuclei were stained with Vectashield Mounting medium with DAPI (Vector, cat#: H1200). The images were captured using an Olympus IX81 fluorescence microscope with SlideBook 6 (Intelligent-Image) software and quantified using ImageJ software (NIH). The apoptotic cells and efferocytosis in atherosclerotic lesions were measured by TUNEL and macrophage staining of cross-sections of the proximal aorta. Briefly, 5 μm sections were fixed in 2% paraformaldehyde, treated with 3% citric acid, and then stained using the in situ TMR red cell death detection kit (Roche Applied Science, cat#: 12156792910) [39]. Macrophages were stained with anti-MOMA2 antibody and nuclei were counterstained using Hoechst. TUNEL-positive (TUNEL+) cells and efferocytosis were then quantitated in proximal aortic sections as previously described [39].

### 2.14. Masson’s Trichrome staining

Masson’s Trichrome staining procedure (Sigma, cat#: HT15-1KT) was used for measurement of lesion necrotic core size and fibrous cap thickness as previously described [36, 39] Briefly, 5 μm sections of the proximal aorta were fixed for 30 min with Bouin’s solution and stained with hematoxylin for nuclei (black). Biebrich scarlet solution and phosphotungstic/phosphomolybdic acid were added to stain cytoplasm (red), and Aniline blue solution was applied to stain collagen (blue). Images were obtained and analyzed for fibrous cap thickness and % necrotic area using ImageJ software (NIH).

### 2.15. Measurement of prostaglandin (PG) metabolites

Male *Ldlr*^-/-^ mice were given vehicle (n=8) or 1 mg/mL PPM (n=8) for 8 weeks. Urine was collected for 6 h in metabolic cages with 2 mice per cage after 8 weeks of treatment with PPM and analyzed by LC/MS for 2,3-dinor-6-ketoPGF1 (2,3DN) and 11-dehydro TxB2 (11dTxB2) by the Eicosanoid Analysis Core at Vanderbilt University. The creatinine levels are measured using a kit (Enzo Life Sciences; cat # ADI-907-030A). The urinary metabolite levels in each sample were normalized using the urinary creatinine level of the sample and expressed as ng/mg creatinine.

### 2.16. Measurements of aortic ONE-Lysyl adducts and IsoLG-lysyl-lactam adducts

The aortic tissues were isolated, homogenized and then the ONE-Lysyl and IsoLG-lysyl adducts were measured by LC/MS/MS using a Waters Xevo-TQ-Smicro triple quadrupole mass spectrometer as previously described [36].

### 2.17. RNA isolation and Quantitative RT-PCR

Total tissue and cell RNA were extracted and purified using PureLink RNA Mini kit (Invitrogen 12183018A) as described in the manufacturer’s protocol. Complementary DNA was synthesized with iScript reverse transcriptase (Bio-Rad). Relative quantitation of the target mRNA was performed using specific primers, SYBR probe (Bio-Rad) and iTaqDNA polymerase (Bio-Rad) on IQ5 Thermocylcer (Bio-Rad). The data were analyzed using the 2^−ΔΔCt^ method, and the results of are presented as the fold change of target gene expression in a sample relative to a reference sample, normalized to a reference gene (18S) [36, 40].

### 2.18. Flow analysis of white blood cells

To measure the numbers of blood neutrophils, Ly6C^hi^ monocytes, and Ly6C^low^ monocytes in *Ldlr*^-/-^ mice, blood cells were collected and stained with a cocktail of antibodies targeting these cells [37]. Briefly, 100 μL of blood was collected using Natelson Blood collection tubes, and then incubated with Fc inhibitor CD16/32 for 10 min at RT. Lysis buffer (3 mL) was pre-warmed at 37°C, added to the blood samples, and incubated at room temperature for 30 min to lyse red blood cells. Samples were centrifuged at 1200 rpm for 10 mins, and the cell pellets were resuspended with 300 ul of FACS buffer (2% FBS in PBS). Antibodies Pacific blue Ly6G (1A8, Biolegend), PE-Cy7 CD11b (M1/70, Biolegend), PE anti-mouse CD3 Antibody (17A2, Biolegend), FITC Anti-mouse CD19 for B cells (6D5, Biolegend) and APC/Cy7 anti-mouse Ly6C (HK1.4, Biolegend) were added, and then incubated with cells on ice for 45 min. The blood T regulatory (Treg) and T helper 2 (Th2) cells were detected using antibodies, where CD3+FoxP3+ positive cells were determined as Treg cells and CD4+GATA-3+ positive populations were Th2 cells. For all flow cytometry assays where propidium iodide or 7-AAD dye were not used, we used gates based on the forward and side scatter to remove dead cells and debris for cell viability from the analysis. Cells were washed with FACS buffer twice and analyzed using FACS DiVa v6.1 software (BD Biosciences) in the Research Flow Cytometry Core Laboratory, Veterans Administration Medical Center.

### 2.19. Bone marrow cell analyses and proliferation of monocytes and HSPC

The bone marrow cells were treated with 5 μm EdU for 24 hours to detect proliferation. The cells were then washed, blocked with Fc inhibitor CD16/32, and stained with FITC anti-mouse CD11b (monocyte marker) or CD34 (HSPC marker, eBioscience, Cat#: 2356471), and EdU positive cells were detected using Click-iT EdU Flow Cytometry Assay Kit (Invitrogen, C10646). Briefly, cells were fixed and permeabilized, and stained with Alexa Fluor 594 picolyl azide in Click-iT EdU buffer following the manufacturer’s protocol. For the fluorescence microscopy assay, the stained cells were placed on glass slides, the images captured, and dual CD11b/EdU positive cells were counted as proliferating HSPC. For the flow cytometry assay, dual CD34/EdU positive cells were considered as proliferating HSPC using FACS DiVa v6.1 software and equipment (BD Biosciences).

### 2.20. Statistics

The data are presented as mean□±□SEM visualized by either box plots or bar charts. The normality of the sample populations was examined by the Kolmogorov-Smirnov test. Between-group differences were assessed with two-sided unpaired t-test (2 groups) and one-way ANOVA (>2 groups, Bonferroni’s correction for multiple comparisons). Their nonparametric counterparts, Mann–Whitney test (2 groups) and nonparametric Kruskal–Wallis test (more than 2 groups, Bunn’s correction for multiple comparison), were used when assumptions for parametric methods were not satisfied. The statistical analysis was performed using Graphpad Prism 7. Statistical tests were determined to be statistically significant at two-sided significance level of 0.05 after correction for multiple comparisons (*P<0.05, **P<0.01, ***P<0.001).

## 3. RESULTS

### 3.1. Effects of PPM on ApoAI crosslinking, HDL cholesterol efflux capacity, and MDA-LDL adducts

As MDA-apoAI adducts have been localized to atherosclerotic lesions [21], and oxidation of HDL lipids using a MPO/H_2_O_2_/Cl^−^ system promotes formation of apoAI-MDA adducts [19], we examined the in vitro effects of PPM on MDA-mediated apoAI crosslinking and impairment of net cholesterol efflux capacity. Modification of HDL with 250 μM MDA increased the degree of apoAI crosslinking (Figures 1A and 1B) and decreased the capacity of the HDL to reduce *Apoe*^-/-^ macrophage cholesterol content by 64% (Figure 1C). Addition of PPM significantly decreased the MDA-mediated crosslinking of apoAI (Figures 1A and 1B) and improved the impairment of HDL net cholesterol efflux capacity (Figure 1C). Besides lipid dicarbonyl modification of apoAI, the MPO generated oxidant, hypochlorous acid (HOCl), can also directly modify apoAI resulting in chlorinated tyrosine or oxidized tryptophan residues, which promote crosslinking and impair cholesterol efflux [13–15]. As PPM is an electron rich molecule that may readily react with HOCl [41], we investigated whether PPM could also inhibit HOCl mediated crosslinking of lipid-free apoAI (Figure 1D). Similar to other studies [13–15], when lipid-free apoAI was incubated with HOCl, there was extensive crosslinking, and consistent with PPM neutralization of HOCl [41], PPM prevented HOCl mediated crosslinking of lipid-free apoAI (Figure 1D). In addition, MPO-mediated HDL modification reduced the net cholesterol efflux capacity by 54.5% and PPM prevented any impairment in HDL cholesterol efflux capacity in *Apoe*^-/-^ macrophages (Figure 1E). More importantly, administration of PPM to *Ldlr*^-/-^ mice on a western diet increased the net cholesterol efflux capacity of HDL (Figure 1F) showing that PPM scavenging in vivo preserves HDL function. As apoB adducts increase foam cell formation and are localized to plaques [42–44], we examined the effects of PPM on LDL MDA content. Compared to treatment with water alone, PPM reduced the MDA-LDL adducts by 40% in *Ldlr*^-/-^ mice on a WD for 16 weeks (Figure 1G) [9, 19].

**Figure 1.**
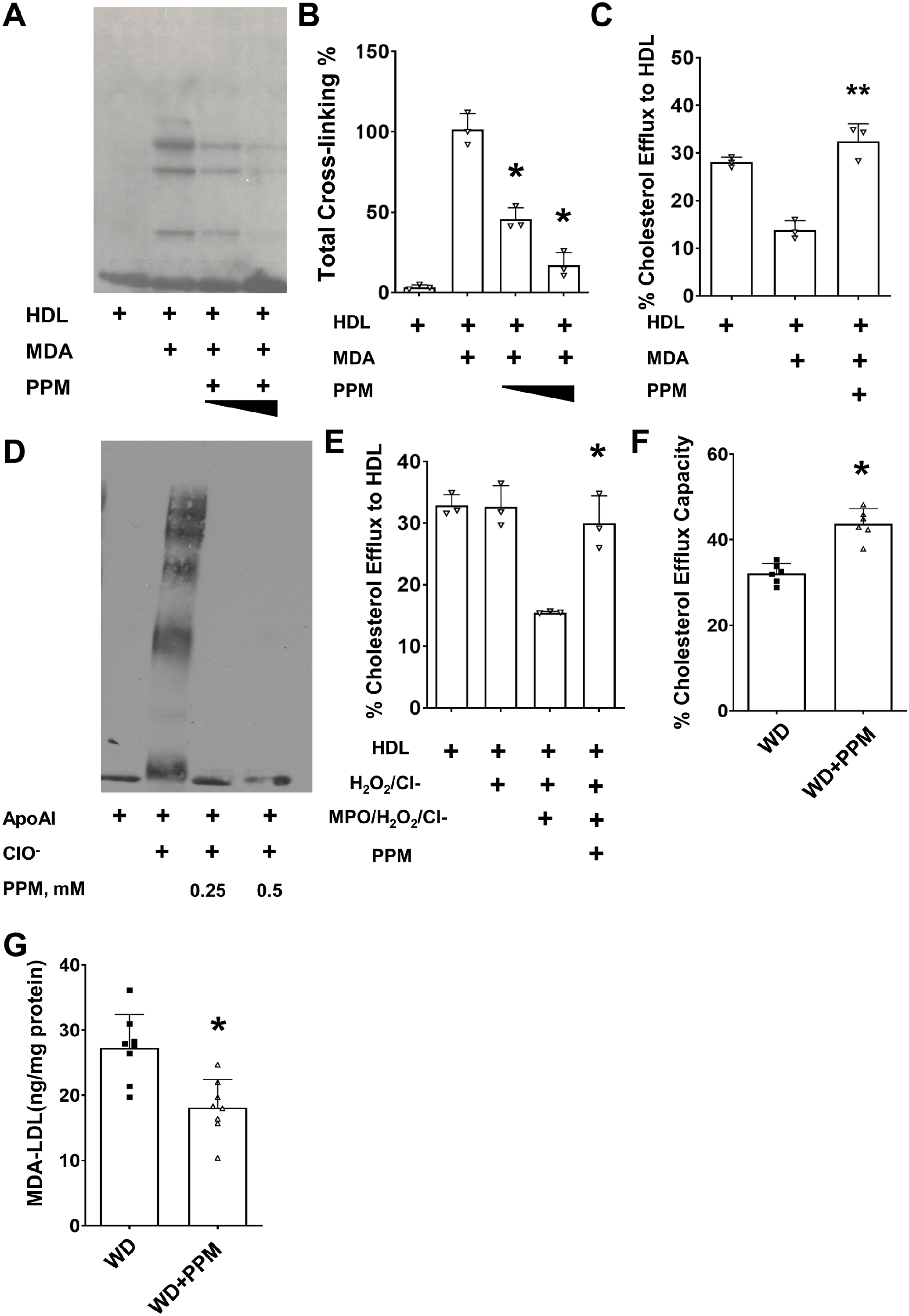
PPM prevents MPO- and MDA-mediated HDL modification and preserves HDL cholesterol efflux capacity. A-B. PPM prevents the crosslinking of apoAI on HDL caused by MDA modification. C. PPM prevents MDA mediated HDL cholesterol efflux dysfunction in *Apoe*^-/-^ macrophages. Cholesterol-enriched *Apoe*^-/-^ macrophages were incubated for 24h with DMEM alone or containing 50μg HDL protein/ml, and the % decrease in cellular cholesterol was then measured as described in the Methods. D. PPM prevents the HOCl mediated crosslinking of lipid-free apoAI. E. PPM prevents MPO-mediated HDL modification and preserves HDL cholesterol efflux capacity in *Apoe*^-/-^ macrophages. Cholesterol-enriched *Apoe*^-/-^ macrophages were incubated for 24h with DMEM alone or containing HDL (50μg protein/ml) and the % decrease in cellular cholesterol was measured. F. PPM treatment improves the cholesterol efflux capacity of HDL in *Ldlr*^-/-^ mice. Cholesterol-enriched *Apoe*^-/-^ macrophages were incubated for 24h with DMEM alone or containing 2.5% (vol/vol) apoB-depleted serum, and the cellular cholesterol was then measured. G. PPM reduces the plasma levels of MDA-LDL in *Ldlr*^-/-^ mice on WD for 16 weeks. Graph data are expressed as mean ± SEM (n=3 in each group); The in vitro studies are 3 independent experiments, *P <0.05, ** P<0.01 by t-test or one-way ANOVA with Bonferroni’s post hoc test.

### 3.2. Effect of PPM on IR and hepatic fat in *Ldlr*^-/-^ mice

As oxidative stress and HDL dysfunction impact IR, we next examined whether PPM impacts insulin sensitivity in male *Ldlr*^-/-^ mice, which develop IR and increased fat deposition when consuming a WD versus a chow diet [45, 46]. After 16 weeks on a WD, male *Ldlr*^-/-^ mice had significantly increased fasting blood glucose and insulin levels versus chow diet (CD) fed controls (Figures 2A and 2B). Administration of PPM to *Ldlr*^-/-^ mice consuming a WD reduced the fasting glucose and insulin levels (Figures 2A and 2B). We also examined HOMA-IR levels as an indicator of IR and found that treatment with PPM reduced the HOMA-IR in *Ldlr*^-/-^ mice on a WD to HOMA-IR values that were similar to those observed in *Ldlr*^-/-^ mice consuming a chow diet (Figure 2C). To determine whether PPM improved insulin or glucose tolerance after 4hrs fasting, GTT and ITT were done after 12 weeks of male *Ldlr*^-/-^ mice consuming a WD and being treated with PPM (Figures 2D-2G). Treatment of *Ldlr*^-/-^ mice consuming a WD with PPM nearly normalized the dynamic change of glucose levels in response to glucose challenge and significantly decreased the glucose tolerance test area under the curve (AUC) (Figures 2D and 2E). Consistent with the GTT results, PPM reduced the dynamic changes of glucose levels in response to insulin challenge in male *Ldlr*^-/-^ mice and significantly reduced the insulin tolerance test AUC (Figures 2F and 2G). We next examined the effects of PPM on hepatic fat accumulation in male *Ldlr*^-/-^ mice. Compared to male *Ldlr*^-/-^ mice on a chow diet, the hepatic fat levels were increased by 3.6-fold in *Ldlr*^-/-^ mice consuming a western diet (Figures 2H and 2I). Consistent with oxidative stress impacting fat deposition, PPM prevented the hepatic fat accumulation in *Ldlr*^-/-^ mice on a western diet (Figures 2H and 2I). Similar to other reports [46, 47], administration of a western versus chow diet to female *Ldlr*^-/-^ mice did not affect fasting glucose levels or the response to glucose challenge, and PPM had no impact on fasting glucose levels and GTT (Figures S1A-S1C). Taken together, these results demonstrate that scavenging reactive dicarbonyls with PPM protects against IR and hepatic fat accumulation in male *Ldlr^-/-^* mice.

**Figure 2.**
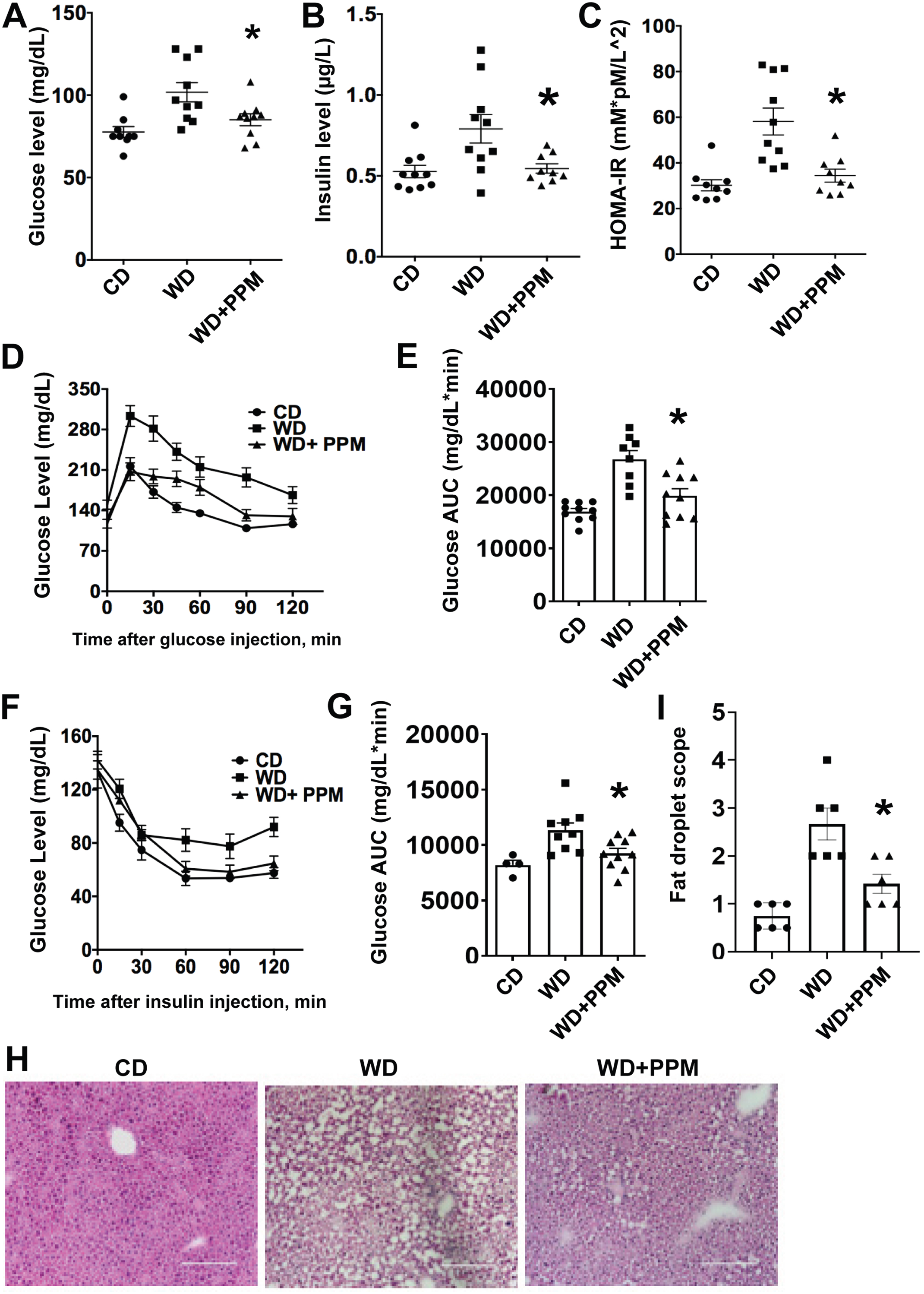
PPM reduces fasting glucose and insulin levels and improves insulin sensitivity in male *Ldlr*^-/-^ mice. A-B. PPM reduces fasting glucose and insulin levels in male *Ldlr*^-/-^ mice. C. PPM normalizes HOMA-IR values in male *Ldlr*^-/-^ mice (n=9 in group on a chow diet, n=10 in group on a western diet, n=9 in group on a western diet and treated with PPM). D-E. Effect of PPM on glucose levels in male *Ldlr*^-/-^ mice in response to injection of glucose using IPGTT as described in Methods (n=9-10 in each group). F-G. Effect of PPM on glucose levels in male *Ldlr*^-/-^ mice in response to injection of insulin using IPITT as detailed in Methods. AUC, area under the curve. H-I. PPM prevents hepatic fat deposition in *Ldlr*^-/-^ mice (n=6), as determined by H&E staining (H) and measured using Image J (I). Scale bar= 100 μm. A.U., arbitrary units. For each experiment, graph data are expressed as mean ± SEM; *P <0.05 by one-way ANOVA with Bonferroni’s post hoc test.

### 3.3. Effects of PPM on atherosclerosis extent and plaque dicarbonyl-lysyl adducts

To examine the effect of scavenging reactive dicarbonyls on atherosclerosis, male *Ldlr*^-/-^ mice were fed a WD for 16 weeks and treated with water alone or containing PPM. As shown in Figures 3A-3B and S2A-S2C, PPM did not change the plasma cholesterol, TG levels, body weight, water intake, or ALT activities. Importantly, administration of PPM versus water alone reduced the extent of proximal aortic atherosclerosis by 46% (Figures 3C-3D). In addition, PPM treatment reduced the en face aortic lesion content by 52% in male *Ldlr*^-/-^ mice (Figures 3E-3F). We next examined the effects of PPM on atherosclerosis in female *Ldlr*^-/-^ mice, which studies suggest are more susceptible to atherosclerosis than male *Ldlr*^-/-^ mice [48]. Similar to male *Ldlr*^-/-^ mice, PPM treatment of female mice on a western diet for 16 weeks reduced the extent of proximal aortic atherosclerosis by 48% without changing plasma cholesterol, triglyceride levels, or lipoprotein FPLC profile (Figures S3A-S3E). As with male *Ldlr*^-/-^ mice, PPM did not change body weight, water intake, and ALT activities in female *Ldlr*^-/-^ mice (Figures S2D-S2F). Consistent with PPM being an effective scavenger of reactive dicarbonyls, immunohistochemistry analysis of MDA-lysyl adducts in plaques revealed that PPM significantly reduced the MDA adduct-positive area in the proximal aortic lesions of female *Ldlr*^-/-^ mice (Figures S4A-S4B). In addition, compared to water alone, treatment with PPM significantly decreased the levels of ONE-lysyl adduct in aortic lesions of male *Ldlr*^-/-^ mice (Figure S4C). PPM also impacted the levels of IsoLG-lysyl adduct in the aorta of male *Ldlr*^-/-^ mice, with a trend towards reduced levels of adducts (p=0.058, Figure S4D). These results demonstrate that PPM treatment effectively decreases atherosclerosis development in experimental mouse models without changing plasma cholesterol and suggests that improved HDL function as well as reduced apoB modification and decreased plaque cellular adducts contribute to the impact on atherosclerosis.

**Figure 3.**
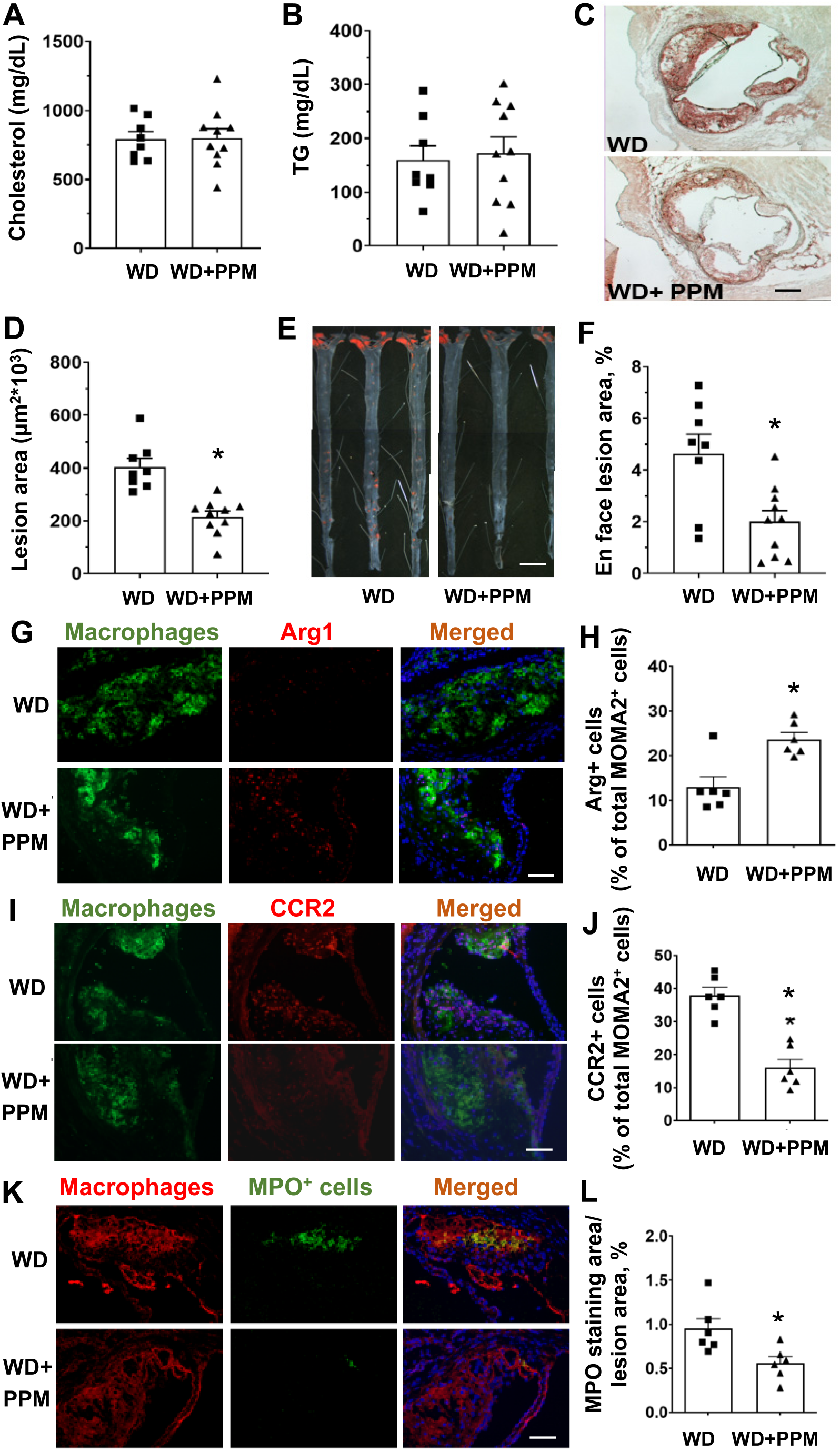
PPM reduces aortic root and en face atherosclerotic lesions in male *Ldlr*^-/-^ mice. A-B. PPM does not affect plasma cholesterol levels or triglyceride levels in male *Ldlr*^-/-^ mice. C-D. PPM reduces the aortic root atherosclerotic lesions in male *Ldlr*^-/-^ mice. E&F. PPM reduces en face atherosclerotic lesion area in male *Ldlr*^-/-^ mice. Male *Ldlr*^-/-^ mice were pretreated with 1 g/L of PPM for 2 weeks on chow diet. Then mice were treated with 1 g/L of PPM for 16 weeks on WD. Oil-Red-O staining of atherosclerosis lesion in male *Ldlr*^-/-^ mice with vehicle (n=8) or PPM (n=10) is shown. Scale bar = 200 μm. Data are expressed as mean ± SEM; *P <0.05, ** P<0.01, ***P<0.001 by two-sided unpaired t-test. G&H. Immunohistochemistry staining of Arg1 + macrophages was performed as described in Methods. H. PPM increases the number of Arg1 + macrophages in lesions of *Ldlr*^-/-^ mice. I&J. Immunohistochemistry staining of CCR2+ macrophages was performed as described in Methods. J. PPM reduces the number of CCR2+ macrophages in lesions of *Ldlr*^-/-^ mice. K&L. Pro-inflammatory enzyme MPO staining of atherosclerotic lesions in *Ldlr*^-/-^ mice is shown. L. PPM reduces MPO expression in atherosclerotic lesions of *Ldlr*^-/-^ mice. G, I, & K. Scale bar = 40 μm. n=6 in each group, for each experiment, graph data are expressed as mean ± SEM; *P <0.05 by two-sided unpaired t-test.

### 3.4. In vivo and in vitro effects of PPM on macrophage inflammation

As macrophage inflammation is modulated by HDL and oxidative stress, we next examined the effects of scavenging reactive dicarbonyls on atherosclerotic lesion macrophage inflammatory markers. Immunohistochemical staining of the proximal aortic sections from male *Ldlr*^-/-^ mice demonstrates that compared to water alone, PPM increased the number of anti-inflammatory Arg1 positive M2 macrophages by 120% (Figures 3G and 3H) and decreased the numbers of pro-inflammatory CCR2 positive macrophages by 58% (Figures 3I and 3J). In addition, PPM treatment reduced the expression of MPO, which is a marker of neutrophil extracellular traps (NETs) that promote the conversion of anti-inflammatory macrophages to a proinflammatory phenotype [49, 50], in the proximal aortic lesions of male *Ldlr*^-/-^ mice (Figures 3K and 3L). In addition, similar effects of PPM on macrophage Arg1 levels were seen in proximal aortas of female *Ldlr*^-/-^ mice (Figures S3F-S3G). Examination of the serum proinflammatory cytokines revealed that scavenging reactive dicarbonyls with PPM decreased the levels of IL-1β and TNF-α by 34% and 62% in male *Ldlr*^-/-^ mice (Figures 5SA-5SB). We also examined the effects of PPM versus water alone on hepatic inflammatory markers in male *Ldlr*^-/-^ mice consuming a western diet for 16 weeks. The mRNA levels of proinflammatory CCL2 and CCL4 were decreased by 20%, and 23% in PPM compared to water alone treated *Ldlr*^-/-^ mice (Figures S5C-S5D). In contrast, PPM increased the hepatic mRNA levels of anti-inflammatory Arg1 by 2.2–fold (Figure S5E). PPM did not significantly impact the hepatic mRNA levels of CCL3 and CXCL2 in male *Ldlr*^-/-^ mice (Data not shown). Importantly, PPM treatment of male *Ldlr*^-/-^ mice reduced the aortic levels of IL-1β and IL-6 mRNA (Figures S6A-S6B). Next, we examined the ex vivo effects of PPM on macrophage inflammatory markers. Thioglycolate elicited peritoneal macrophages were isolated from PPM and water alone treated male *Ldlr*^-/-^ mice consuming a WD for 16 weeks and then stimulated with oxidized LDL or LPS. Macrophages from PPM versus vehicle treated *Ldlr*^-/-^ mice had 28% and 33% reductions in mRNA levels of IL-1β and IL-6 in response to oxidized LDL (Figures 4A-4B). In addition, there was a significant reduction in expression of IL-1β and IL-6 in LPS stimulated macrophages from PPM versus vehicle treated *Ldlr*^-/-^ mice (Figures 4C-4D). In contrast, PPM versus water alone administration increased Arg1 expression by 3.6-fold in LPS stimulated macrophages (Figure 4E). Importantly, in vitro incubation of macrophages with PPM and oxidized LDL compared to oxidized LDL alone markedly reduced the mRNA levels of IL-1β and TNF-α (Figures 4F-4G) suggesting that PPM minimizes inflammation by preventing cellular oxidative modifications. Due to the striking impact on inflammatory cytokines and macrophage phenotype in vivo, we also measured urinary prostaglandins to evaluate whether PPM might be inhibiting cyclooxygenase (COX). Urine samples were analyzed for 2,3-dinor-6-ketoPGF1 (2,3DN) and 11-dehydro TxB2 (11dTxB2) by LC/MS/MS. We found that there were no significant differences in levels of these major urinary prostaglandin metabolites of *Ldlr*^-/-^ mice treated with PPM compared to the vehicle control (Figure S7), indicating that PPM was not significantly inhibiting COX in vivo in mice.

**Figure 4.**
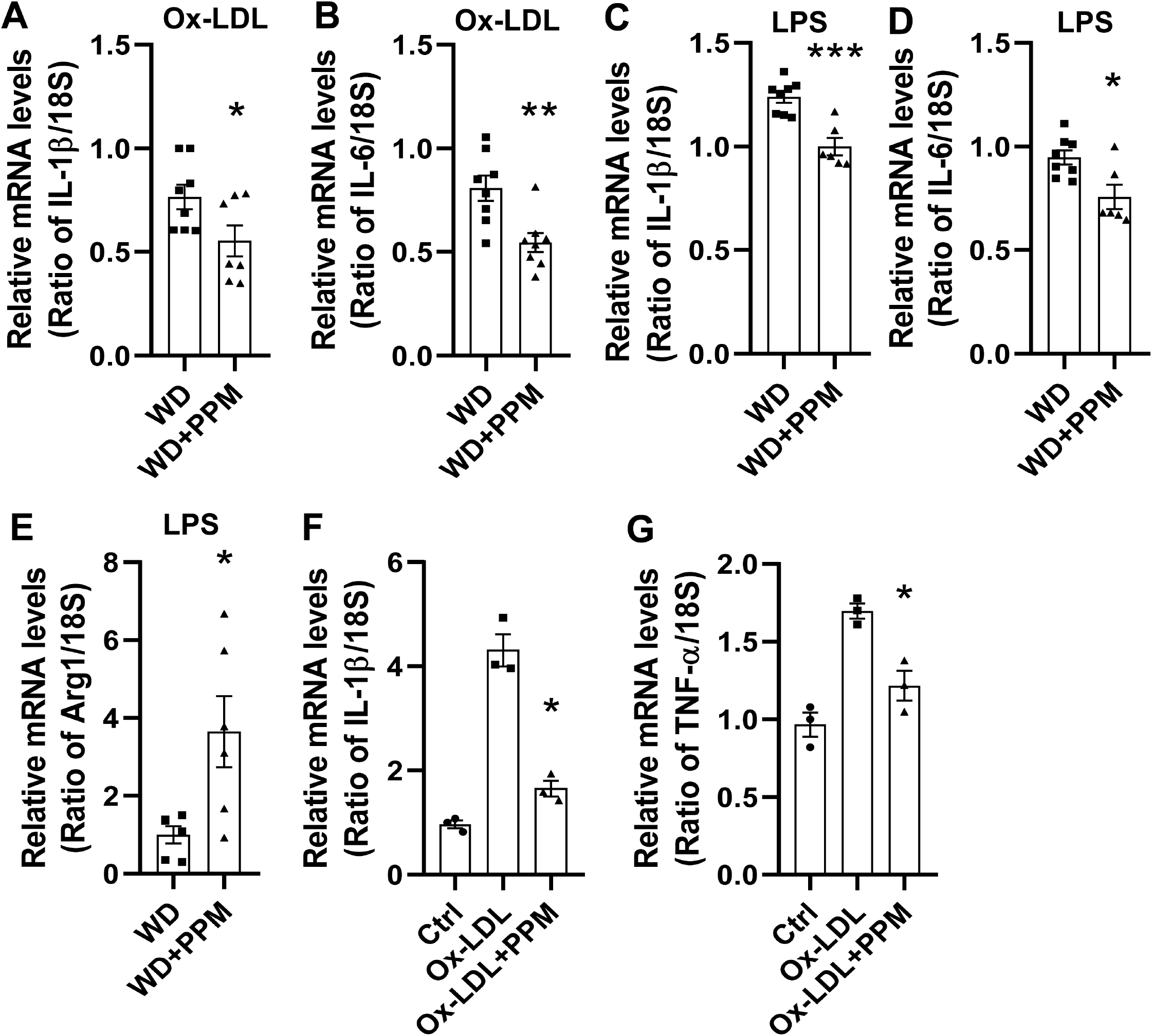
PPM reduces the macrophage inflammatory response to oxidized LDL and LPS. A-E. The ex vivo effect of PPM on the expression of inflammatory genes in macrophages incubated with either oxidized LDL (A-B) or LPS (C-E). A-E. Macrophages were isolated from male *Ldlr*^-/-^ mice that were fed a western diet for 16 weeks and were treated with vehicle alone or with 1 g/L of PPM. The cells were then incubated for 24h with 40 μg/mL of oxidized LDL (A-B) or 4h with 100 ng/ml of LPS (C-E). A-B. In vivo treatment with PPM reduces the mRNA levels of IL-1β and IL-6 in *Ldlr*^-/-^ macrophages stimulated with oxidized LDL. C-E. PPM reduces the ex vivo expression of IL-1β and IL-6 and increases Arg1 mRNA levels in *Ldlr*^-/-^ macrophages in response to LPS. A-E. The mRNA levels were measured by real-time PCR as described in Methods. n=7 in each group, for each experiment, graph data are expressed as mean ± SEM; *P <0.05 by two-sided unpaired t-test. F-G. PPM reduces the mRNA expression levels of IL-1β and TNF-α in *Apoe*^-/-^ macrophages in response to 24h with 40 μg/mL ox-LDL. n=3 independent experiments. Data are expressed as mean ± SEM; *P <0.05, **P <0.01, by two-sided unpaired t-test.

**Figure 5.**
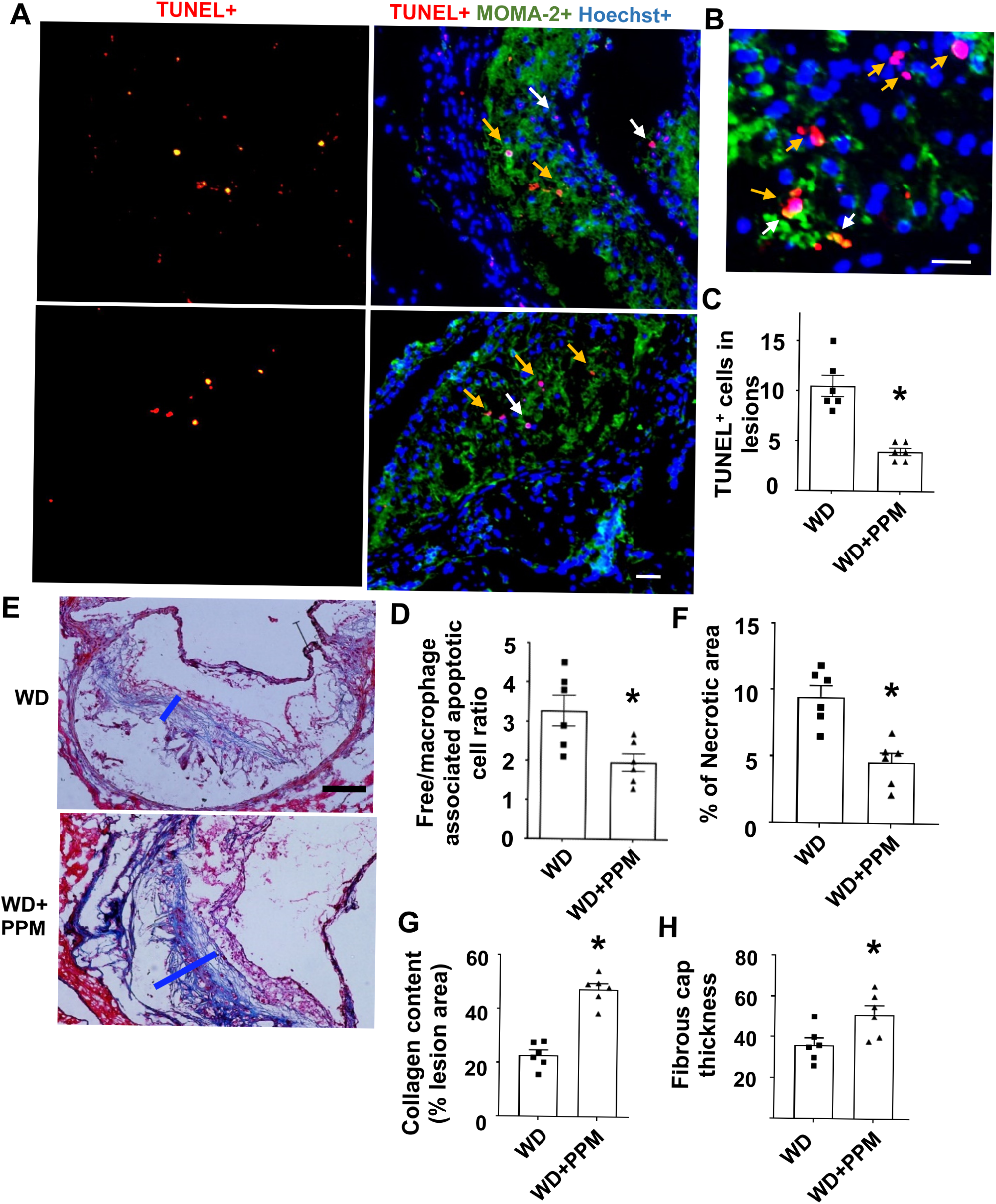
PPM reduces the number of apoptotic cells, increases efferocytosis, and stabilizes the atherosclerotic plaque of male *Ldlr*^-/-^ mice. A-B. Images depict TUNEL staining of nuclei (red) and merged images show TUNEL+, MOMA2+ (green) and Hoechst+ (blue) staining of atherosclerotic lesions in male *Ldlr*^-/-^ mice. Scale bar = 50 μm. C. PPM reduces the apoptotic cells in atherosclerotic lesions of male *Ldlr*^-/-^ mice (n=6). B&D. PPM enhances macrophage efferocytosis of apoptotic cells. Higher magnification images of the TUNEL staining (B) were used to quantitate (D) efferocytosis of dead cells in aortic root sections (n=6 per group). Efferocytosis was quantitated as the free (white arrows) vs macrophage-associated TUNEL-positive cells (yellow arrows) in the proximal aortic sections (D). E-H. The plaque stability was measured by using Trichrome staining of atherosclerotic lesions of *Ldlr*^-/-^ mice (n=6 in each group). The necrotic area (F), collagen content (G), and fibrous cap thickness (H) and necrotic area were quantitated in atherosclerotic lesions of *Ldlr*^-/-^ mice (n=6 in each group). Scale bar = 40 μm. For each experiment, graph data are expressed as mean ± SEM; *P <0.05 by two-sided unpaired t-test.

### 3.5. Effects of PPM on plaque cell death, efferocytosis, and features of stability

As plaque inflammation and oxidative stress induces cellular death, we next investigated the effects of PPM on lesion cell death. PPM versus vehicle treatment decreased the number of dead macrophages as evidenced by TUNEL staining in the proximal aortic lesions in both male and female *Ldlr*^-/-^ mice (Figures 5A-5C and Figures S8A-S8B). From the costaining of TUNEL and live macrophages, the effect of PPM on efferocytosis of apoptotic cells in male and female *Ldlr*^-/-^ mouse plaques was also examined (Figures 5A, 5B, and 5D and Figures S8A and S8C). The number of TUNEL-positive cells not associated with macrophages was decreased by 40% in lesions of male mice treated with PPM versus vehicle (Figure 5D). Similar results on the effects of PPM on plaque efferocytosis were observed in female *Ldlr*^-/-^ mice (Figures S8A and S8C). Consistent with the decreased inflammatory cell death and enhanced efferocytosis, PPM versus vehicle treatment decreased the necrotic area by 51.6% and increased the collagen content and fibrous cap thickness by 107.3% and 41.9% in male *Ldlr*^-/-^ mice, respectively (Figures 5E-5H).

### 3.6. Effects of PPM on plaque T and B cells

As macrophage inflammation and efferocytosis are impacted by T cell populations, we next examined the effects of PPM on plaque T cells. There were similar total numbers of T cells (CD3+) in plaques of PPM versus control treated female *Ldlr*^-/-^ mice consuming a western diet for 14 weeks (Figures 6A-6B). Interestingly, the number of Th2 cells (GATA-3+) were increased by 2.2-fold in plaques of *Ldlr*^-/-^ mice (Figures 6C-6D). In addition, the number of Tregs (Fox3P+) which promote efferocytosis and reduce inflammation were increased by 2.1–fold in PPM versus control *Ldlr*^-/-^ mice (Figures 6E-6F). PPM did not affect the number of B cells (B220+) in plaques of *Ldlr*^-/-^ mice (Figures S9A-S9B). Taken together, our data show that PPM suppresses the features of vulnerable plaques in a hypercholesterolemic murine model.

**Figure 6.**
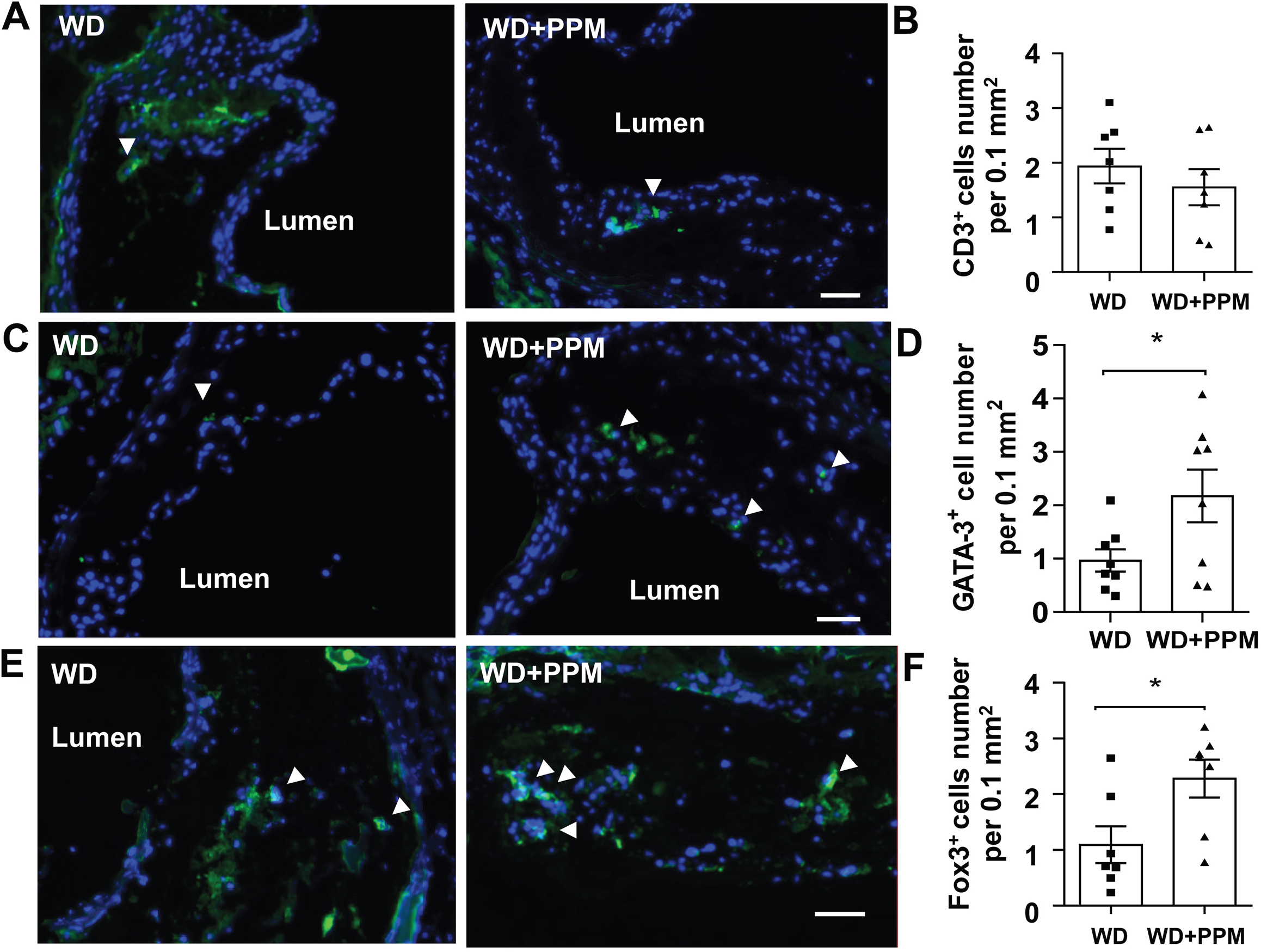
PPM increases the number of Treg and Th2 lymphocytes in plaques of *Ldlr*^-/-^ mice. A-B. Immunohistochemistry staining and quantitation of CD3+ (green) T lymphocytes in atherosclerotic lesions of female *Ldlr*^-/-^ mice. Scale bar = 50 μm. C-D. Immunohistochemistry staining and quantitation of GATA-3+ (green) Th2 lymphocytes in atherosclerotic lesions of female *Ldlr*^-/-^ mice. Scale bar = 50 μm. E-F. Immunohistochemistry staining of FoxP3+ (green) Treg cells. F. PPM increases the number of Treg cells in plaques of *Ldlr*^-/-^ mice. Scale bar = 50 μm. For each experiment (n=6 or 7 in each group), graph data are expressed as mean ± SEM; *P <0.05 by two-sided unpaired t-test.

### 3.7. Effects of PPM on white blood cell populations in *Ldlr*^-/-^ mice

Studies have demonstrated that Ly6C^hi^ monocytes preferentially infiltrate the arterial wall and convert to inflammatory M1-like macrophages during inflammation thereby promoting progression of the atheroma [37, 51, 52]. As inflammatory Ly6C^hi^ monocytosis is reduced by HDL cholesterol efflux and enhanced by oxidative stress, we next examined the effects of reactive dicarbonyl scavenging on subsets of white blood cells in female *Ldlr*^-/-^ mice consuming a western diet for 14 weeks (Figures 7A-7B). Interestingly, we found that compared to water alone, PPM treatment of female *Ldlr*^-/-^ mice decreased the total number of blood monocytes and markedly reduced the number of Ly6C^hi^ monocytes (Figures 7A-7D). In addition, PPM did not affect the number of blood Ly6C^low^ cells but decreased the ratio of Ly6C^hi^ to Ly6C^low^ monocytes (Figures 7E-7F) in female *Ldlr*^-/-^ mice. A similar impact of PPM treatment on total monocytes, Ly6C^hi^ cells and Ly6C^low^ cells was observed in male *Ldlr*^-/-^ mice (Figures S10A-S10D). In both female and male *Ldlr*^-/-^ mice, PPM treatment did not impact the number of circulating neutrophils, T cells, and B cells (Figures S11A-S11F). Similar to the atherosclerotic lesions, the number of anti-inflammatory Th2 (GATA-3+) and Treg (Fox3P+) cells were increased in blood from PPM versus water alone treated female *Ldlr*^-/-^ mice (Figures S12A-S12C).

**Figure 7.**
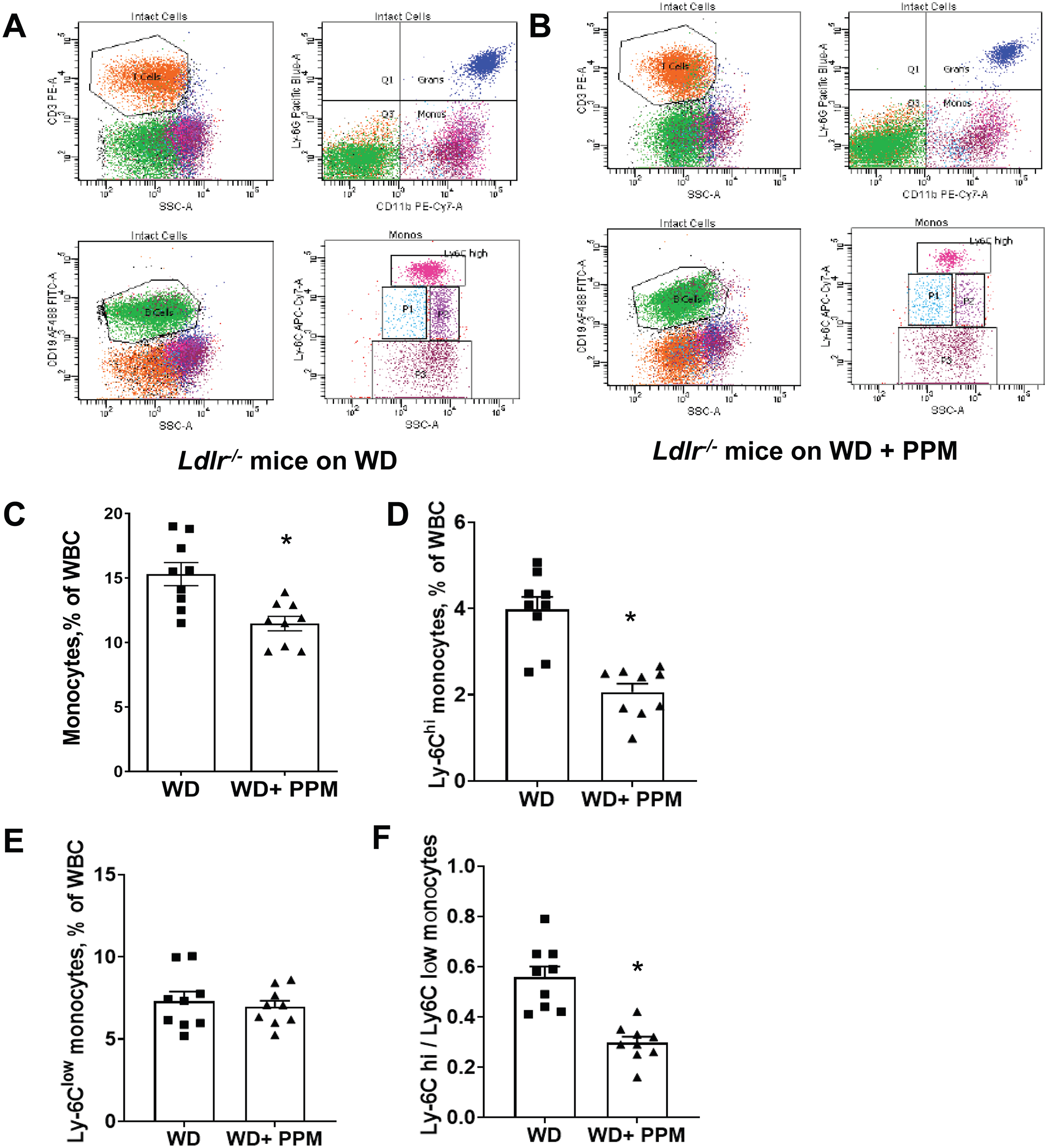
PPM decreases the number of blood Ly6C^hi^ monocytes and the ratio of Ly6C^hi^ to Ly6c^Low^ monocytes in female *Ldlr*^-/-^ mice. A-B. The representative flow cytometry gating strategies for monocyte subsets, neutrophils, T cells, and B cells in peripheral blood of vehicle alone or PPM treated female *Ldlr*^-/-^ mice fed a western diet for 14 weeks are shown. C. There was a reduction in total blood monocytes in PPM versus water treated *Ldlr*^-/-^ mice. D. PPM reduced the number of Ly6C^hi^ monocytes E. PPM did not impact the number of Ly6C ^low^ monocytes. F. PPM decreased the ratio of Ly6C^hi^ to Ly6C ^low^ monocytes. n=9 in each group. Graph data are expressed as mean ± SEM; *P <0.05 by two-sided unpaired t-test.

### 3.8. Effects of PPM on monocyte and HSPC proliferation in *Ldlr*^-/-^ mice

We next examined the impact of PPM on bone marrow Ly6C^hi^ monocytes. Compared to male and female *Ldlr*^-/-^ mice on a chow diet, the numbers of bone marrow Ly6C^hi^ monocytes increased in male and female *Ldlr*^-/-^ mice fed a western diet for 16 weeks (Figures 8A-8B). Consistent with the decreased numbers of blood Ly6C^hi^ monocytes in PPM treated *Ldlr*^-/-^ mice, the number of bone marrow Ly6C^hi^ monocytes trended higher in water alone versus PPM treated male and female *Ldlr*^-/-^ mice fed a western diet (Figures 8A-8B). When the Ly6C^hi^ monocyte data for male and female *Ldlr*^-/-^ mice were grouped together, the number of bone marrow Ly6C^hi^ monocytes was significantly reduced in PPM versus water alone treated *Ldlr*^-/-^ mice (Figure 8C). Consistent with studies demonstrating that oxidative stress enhances inflammatory monocytosis [53], the number of Ly6C^hi^ monocytes were markedly increased in white blood cells incubated with H_2_O_2_ compared with medium alone (Figure 8D). PPM prevented the H_2_O_2_ induced increase in Ly6C^hi^ monocytes (Figure 8D). Examination of the proliferation of monocytes, by Edu incorporation and fluorescence microscopy, in bone marrow cells isolated from male *Ldlr*^-/-^ mice fed a western versus chow diet revealed increased proliferation of CD11b+ monocytes after administration of a western diet (Figures 8E-8F). Importantly, PPM treatment of male *Ldlr*^-/-^ mice decreased the western diet induced increase in proliferation of bone marrow monocytes (Figures 8E-8F). Similar results were observed using fluorescence microscopy to examine the effects of PPM on monocyte proliferation in bone marrow cells isolated from female *Ldlr*^-/-^ mice fed a western diet (Figures S13A-S13B). In addition, measurement of monocyte proliferation as measured by Edu incorporation into CD11b+ cells and flow cytometry revealed similar results with bone marrow isolated from female *Ldlr*^-/-^ mice (Figures S13C-S13D). Next, we examined the effects of PPM on proliferation of CD34+ hematopoietic stem and progenitor cells. Compared to the proliferation of CD34+ HSPC in bone marrow isolated from chow fed male *Ldlr*^-/-^ mice, HSPC proliferation was enhanced in bone marrow from mice consuming a western diet (Figures 8G-8H). Administration of PPM to the male *Ldlr*^-/-^ mice prevented the increase in proliferation of bone marrow CD34+ HSPC (Figures 8G-8H). Similar effects with PPM treatment on the proliferation of CD34+ HSPC were seen in bone marrow isolated from female *Ldlr*^-/-^ mice (Figures S13E-S13F). Collectively, we demonstrate that scavenging reactive dicarbonyls improves HDL function, modulates macrophage polarization, represses Ly6C^hi^ monocytosis, and decreases lesion macrophage inflammation and death resulting in increased plaque stability.

**Figure 8.**
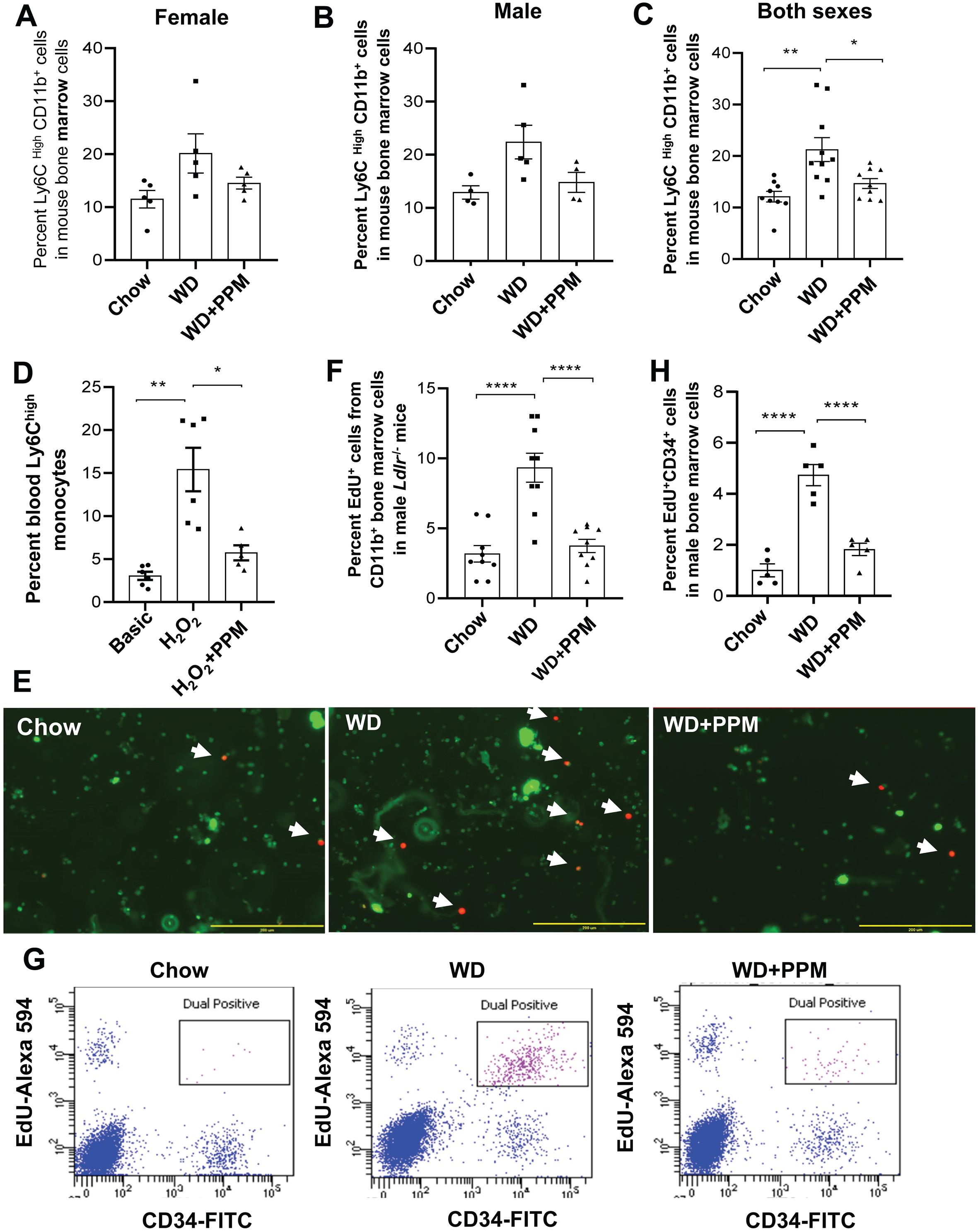
PPM decreases bone marrow Ly6C^hi^ monocytes, oxidative stress induced accumulation of blood Ly6C^hi^ monocytes, and the proliferation of bone marrow monocytes and HSPC. A-C. PPM decreases the number of bone marrow Ly6C^hi^ monocytes. Male (A) or female (B) *Ldlr*^-/-^ mice were fed a chow or a western diet for 16 weeks and treated with vehicle alone or PPM. The bone marrow was isolated and the number of Ly6C^hi^ monocytes was measured by flow cytometry as described in the Methods. C. The data for male (n=4 or 5 per group) and female (n=9 per group) bone marrow Ly6C^hi^ monocytes were grouped. Data are expressed as mean ± SEM; *P <0.05, ** P<0.01 by one-way ANOVA with Bonferroni’s post hoc test. D. PPM decreases the accumulation of Ly6C^hi^ monocytes in white blood cells subjected to oxidative stress. White blood cells isolated from *Ldlr*^-/-^ mice fed a chow diet were incubated for 24h in medium alone or with either H_2_O_2_ (100μM) or H_2_O_2_ and PPM (100 μM). The number of Ly6C^hi^ monocytes was then measured by flow cytometry as described in the Methods. n=6 per group. Data are expressed as mean ± SEM; *P <0.05, ** P<0.01 by one-way ANOVA with Bonferroni’s post hoc test. E-H. PPM decreases the proliferation of bone marrow monocytes (E-F) and HSPC (G-H). Bone marrow was isolated from male *Ldlr*^-/-^ mice fed a chow or western diet for 16 weeks and treated with water alone or with PPM. E-F. Bone marrow cells were incubated for 24h with 5 μM EdU, and then stained with FITC-labeled CD11b antibody (green). EdU (red) was detected as described in the Methods. Dual EdU+ CD11b+ cells were detected by fluorescence microscopy (E) and quantitated (F). n=8 per group. Bar = 200 μM. Data are expressed as mean ± SEM; **** P<0.0001 by one-way ANOVA with Bonferroni’s post hoc test. G-H. Bone marrow cells were incubated for 24h with 5 μM EdU. EdU was detected with Alexa Fluor 594 picolyl azide as described in the Methods. HSPC were stained with FITC-labeled CD34 antibody. Data are expressed as mean ± SEM, N=5 each group, ****P <0.0001 by one-way ANOVA with Bonferroni’s post hoc test.

## 4. DISCUSSION

Oxidative stress plays critical roles in the pathogenesis and progression of atherosclerosis. While some studies administering antioxidants to humans showed decreased atherosclerosis, the majority of clinical trials demonstrated no beneficial effect on cardiovascular events, which may be due to the doses used [54] or from impaired physiologic signaling induced by reactive oxygen species (ROS) that could be atheroprotective. Lipid peroxidation produces reactive dicarbonyls, which covalently bind plasma/cellular proteins, phospholipids, and DNA resulting in altered function and toxicity [55]. We demonstrate that reactive dicarbonyl scavenging with PPM effectively increased HDL cholesterol efflux capacity, (Figures 1 and 2), and reduced IR and atherosclerosis in hypercholesterolemic *Ldlr*^-/-^ mice (Figures 3 and S3) without impacting plasma lipids. In addition, PPM decreased monocytosis, systemic inflammation, and features of plaque instability (Figures 3, 5, 7, 8, S5, S6, S8, S10, and S13). Thus, reactive dicarbonyl scavenging may have therapeutic potential to reduce risk of atherosclerotic clinical events.

PPM reduced atherosclerosis without changing plasma cholesterol (Figures 3 and S3) suggesting that PPM preserves normal molecular functions by preventing oxidative modifications. Consistent with this concept, PPM decreased MDA-, ONE-, and IsoLG-lysyl adducts in plaques of *Ldlr*^-/-^ mice (Figure S4). MPO and dicarbonyl modified LDL enhance foam cell formation and are present in human plaques, [42–44, 56], and PPM reduced plasma MDA-LDL implicating that the atheroprotection is in part due to decreased LDL modification (Figure 1G). Indeed, antibodies against MDA-apoB protect against atherosclerosis development in mice [44]. HDL protects against atherosclerosis via a number of functions including promotion of cholesterol efflux. As oxidative modifications of HDL enhance particle internalization via scavenger receptors [57, 58], we examined the impact of PPM on the ability of HDL to reduce cholesterol mass in cholesterol-enriched macrophages to control for differences in the influx of cholesterol. The removal of cholesterol mass was reduced with MDA and MPO modification of HDL (Figures 1C and 1E), and PPM prevented these impairments in net cholesterol efflux (Figures 1A and 1D). Importantly, in vivo treatment of *Ldlr*^-/-^ mice with PPM increased the HDL net cholesterol efflux capacity (Figure 1F), which probably results from decreased dicarbonyl-ApoAI adducts as we previously showed that PPM treatment of *Ldlr*^-/-^ mice decreases plasma MDA-HDL [19]. In addition, IsoLG or MDA modification of HDL reduces cholesterol efflux capacity in vitro, and HDL from subjects with FH has impaired cholesterol efflux capacity and contains more IsoLG- and MDA adducts compared to HDL from normocholesterolemic controls [19, 20, 22, 36]. HOCl modification of ApoAI (i.e. oxidized tryptophan, chlorinated tyrosine) also reduces cholesterol efflux [13–15], and our studies suggest that PPM decreases HOCl modification of apoAI in vivo as PPM prevented the HOCl-mediated crosslinking of lipid-free apoAI and impairment of cholesterol efflux (Figures 1D-1E). In agreement, the scavenger, pyridoxamine, neutralizes HOCl in vitro and prevents oxidation and chlorination of collagen tryptophan in vivo [41]. It is worth noting that MDA-apoAI and HOCl modified apoAI are present in human plaques [13–15, 21]. Furthermore, HDL cholesterol efflux capacity is independently associated with atherosclerosis, and impairments in HDL cholesterol efflux capacity increase the risk of cardiovascular events [5–7]. Thus, the finding that PPM treatment of *Ldlr*^-/-^ mice enhances HDL net cholesterol efflux capacity highlights the therapeutic potential of in vivo dicarbonyl/oxidant scavenging against atherosclerosis.

Ly6C^hi^ monocytes dominate hypercholesterolemia-associated monocytosis in mice and give rise to inflammatory macrophages in the atheroma [51, 59]. In addition, humans with hypercholesterolemia and coronary artery disease have increased proinflammatory CD14^++^CD16^+^ (intermediate) and CD14^++^CD16^-^ (classical) monocytes, which promote plaque instability and are the human counterparts to mouse proinflammatory Ly6C^hi^ monocytes) [60–64]. Our results show that PPM reduced Ly6C^hi^ monocytosis in response to hypercholesterolemia, providing a novel mechanism whereby dicarbonyl and/or oxidant neutralization decreases atherosclerosis and plaque destabilization (Figures 7 and S10). The effects of PPM on monocytosis resulted from decreased proliferation of bone marrow HSPC and monocytes (Figures 8 and S13), which is probably due in part to the prevention of cellular oxidative modifications. In agreement, PPM prevented in vitro the H_2_O_2_ induced increase in blood Ly6C^hi^ monocytes (Figure 8D). In addition, studies have shown that hypercholesterolemia induces oxidative stress causing enhanced p38 and Notch1 signaling and HSPC proliferation [53, 65]. Furthermore, treatment of hypercholesterolemic *Apoe*^-/-^ and *Scarb1*^-/-^ mice with antioxidants decreases monocytosis [53, 65]. The prevention of lipoprotein modifications likely contributes to the PPM effects on monocytosis. Oxidized LDL enhances HSPC monocyte potential [66]. In addition, HDL reduces proliferation by decreasing ROS in HSPC [53], and, in contrast, HDL dicarbonyl adducts enhance ROS production [67]. Importantly, HDL cholesterol efflux capacity limits HSPC/monocyte proliferation, and PPM improved the ability of HDL from *Ldlr*^-/-^ mice to remove cholesterol (Figure 1F) [68].

Inflammation induces the conversion of infiltrating monocytes into proinflammatory M1-like macrophages, which promote plaque progression and instability [69]. The CANTOS trial supports the importance of inflammation in atherosclerotic cardiovascular disease as treatment of humans with an IL-1β monoclonal antibody lowered recurrent cardiovascular events independent of cholesterol lowering [70]. Our studies show that PPM limits inflammation without impacting plasma cholesterol. Indeed, PPM reduced systemic inflammation as evidenced by reduced serum IL-1β and TNF-α and decreased hepatic inflammatory gene expression in *Ldlr*^-/-^ mice (Figure 4). In addition, PPM reduced plaque M1-like macrophages as substantiated by decreased CCR2 [71] and IL-1β expression (Figure 3 and Figures S3 and S5). Importantly, PPM decreased the association of NETs MPO with plaque macrophages, which propagates oxidative stress and conversion to proinflammatory macrophages [49, 50]. In addition, PPM increased lesional Treg and Th2 cells (Figure 6), which promote conversion to anti-inflammatory M2-like macrophages [72, 73]. In agreement with the increase in M2-like macrophages, the plaques of PPM treated *Ldlr^-/-^* mice contained more Arg1+ macrophages (Figures 3 and S3). In addition, some studies have shown that Tregs enhance efferocytosis signaling via stimulation of macrophage IL-10 production, and Arg1 metabolizes apoptotic cell arginine to promote continual efferocytosis [73, 74]. Consistent with these studies and the increased Tregs and Arg1+ cells, scavenging with PPM enhanced the efferocytosis of lesion apoptotic cells and decreased necrosis (Figures 5 and S8). These anti-inflammatory effects of PPM are in part due to prevention of cellular oxidative modifications as PPM decreased the ex vivo and in vitro inflammatory response to LPS and/or oxidized LDL (Figure S6). In addition, we previously showed that the scavenger, 2-hydroxybenzylamine, decreases the inflammatory response to oxidized LDL and H_2_O_2_ by forming adducts with macrophage dicarbonyls [36]. The prevention of LDL modifications also likely contributed to the PPM effects on plaque macrophage inflammation [56, 75]. Furthermore, functional HDL promotes conversion to anti-inflammatory macrophages by increasing STAT6 mediated expression of Arg1 and Fizz-1 and by limiting inflammatory signaling via ERK1/2 and STAT3 [76, 77]. In contrast, MDA, IsoLG, and HOCl mediated modifications convert HDL into proinflammatory particles [13, 14, 19, 20]. It is also worth noting that HDL anti-inflammatory signaling is intimately linked to cholesterol efflux capacity, [78–80] which is enhanced by PPM scavenging in vivo (Figure 1F).

Similar to other studies, administration of a western diet to male *Ldlr^-/-^* mice caused IR, but had no effect in female *Ldlr^-/-^* mice (Figures 2 and S1) [45–47]. We show that reactive dicarbonyl scavenging decreased IR in male *Ldlr^-/-^* mice, and, consistent with this observation, hepatic fat and inflammation were also reduced (Figures 2 and 4). The effects of PPM occurred without changes in plasma lipids, which substantiates the role of oxidative stress in IR development [81], and is in agreement with studies showing that cellular dicarbonyl adducts impair insulin signaling and glucose uptake in adipocytes, hepatocytes, and muscle cells [82–84]. The preservation of HDL function may also contribute to the decreased IR as the cholesterol transporter, ABCA1, enhances insulin signaling and glucose uptake in adipose and muscle [85, 86]. In addition, glucose metabolism is improved with injection of apoAI, apoAI peptides, or HDL in diabetic mice and humans with T2DM [11, 12, 87]. A weakness of our studies is that Ldlr^-/-^ mice may not be the most stringent model for testing the effects of PPM on IR. However, it is worth noting that our results are substantiated by other studies demonstrating that the parent molecule of PPM, pyridoxamine, improves insulin sensitivity and glucose metabolism in more robust models including obese *db/db* mice and diabetic rats [38, 88].

In summary, dicarbonyl scavenging with PPM decreased atherosclerosis and IR without changing plasma lipid levels. Importantly, PPM reduced monocytosis and enhanced features of plaque stability. The effects of PPM on atherosclerosis and insulin sensitivity are likely due to prevention of oxidative damage to cellular and plasma constituents. In particular, reduced modification of apoB containing lipoproteins and improved HDL cholesterol efflux capacity are atheroprotective. Thus, PPM offers therapeutic potential for addressing the residual cardiovascular risk that persists in patients treated with current LDL-C lowering medications.

## ACKNOWLEDGMENTS

This work was supported by National Institutes of Health Grants: HL116263, HL148137, and HL146134. Jiansheng Huang was supported by America Heart Association Postdoctoral Fellowship (19POST34450266). The analyses of urinary prostaglandin metabolites were performed in the Vanderbilt University Eicosanoid Core Laboratory.

## AUTHOR CONTRIBUTIONS

Jiansheng Huang: Conceptualization, Investigation, Visualization, Writing - Original Draft, Formal analysis; Huan Tao: Investigation, Visualization, Supervision, Writing - Review & Editing, Patricia G. Yancey: Conceptualization, Supervision, Visualization, Formal analysis; Writing - Original Draft; Zoe Leuthner: Investigation, Writing - Review & Editing, Linda S. May-Zhang: Investigation, Visualization, Writing - Review & Editing, Ju-Yang Jung: Investigation, Writing - Review & Editing, Youmin Zhang: Investigation; Lei Ding: Investigation; Venkataraman Amarnath: Resources, Dianxin Liu; Investigation; Sheila Collins: Supervision, Writing - Review & Editing; Sean S. Davies: Supervision, Methodology, Writing - Review & Editing; MacRae F. Linton: Conceptualization, Supervision, Writing - Review & Editing, Formal analysis, Funding acquisition

## DATA AVAILABILITY

The data that support the findings of this study are available from the corresponding author upon reasonable request.

## APPENDIX A. FIGURES A.1 – A.8

## APPENDIX B. SUPPLEMENTARY DATA

Supplementary data to this article can be found at

## Abbreviations

AUC: Area under the curve
Arg1: Arginase1
CCR2: C-C chemokine receptor 2
CD: Chow diet
COX: Cyclooxygenase
EdU: 5-ethynyl-2’-deoxyuridine
FH: Familial hypercholesterolemia
GTT: Glucose tolerance test
HSPC: Hematopoietic stem and progenitor cells
H&E: Hematoxylin and eosin
HDL: High density lipoprotein
HOMA-IR: Homeostatic model assessment for insulin resistance
HOCl: Hypochlorous acid
IR: Insulin resistance
ITT: Insulin tolerance test
IsoLG: Isolevuglandins
LDL: Low density lipoprotein
MDA: Malondialdehyde
MPO: Myeloperoxidase
NETs: neutrophil extracellular traps
PPM: 5’-O-pentyl-pyridoxamine
ONE: 4-Oxo-nonenal
ROS: Reactive oxygen species
RCT: Reverse cholesterol transport
T2D: Type 2 diabetes
WD: Western diet

---

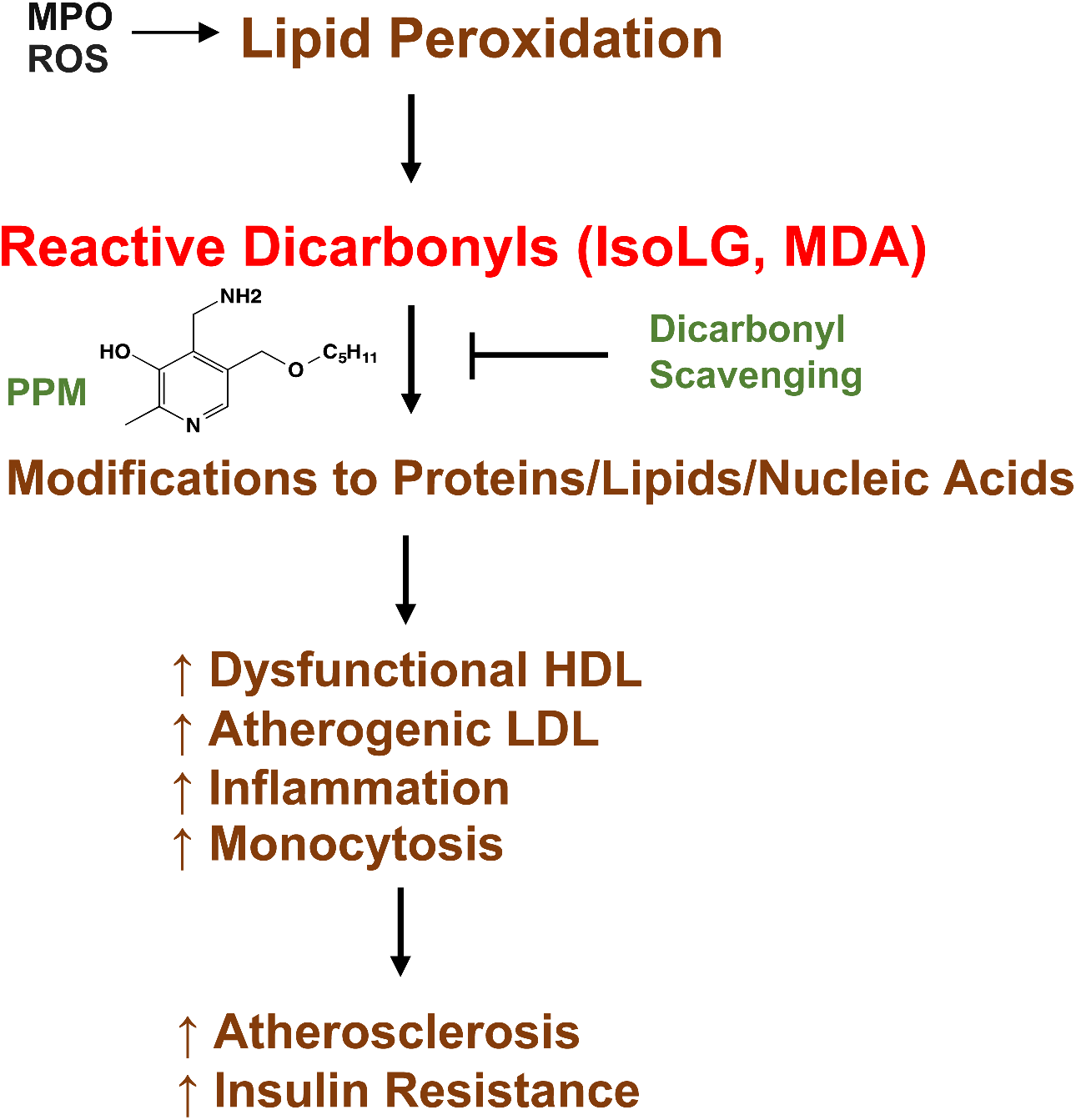

## References

[1] Linton, M.R.F., P.G. Yancey, S.S. Davies, W.G. Jerome, E.F. Linton, W.L. Song, et al., 2019. The Role of Lipids and Lipoproteins in Atherosclerosis. In: K.R. Feingold, et al., Editors. Endotext, South Dartmouth, MA; Available from: https://www.ncbi.nlm.nih.gov/pubmed/26844337.

[2] Sampson, U.K., S. Fazio, and M.F. Linton, 2012. Residual cardiovascular risk despite optimal LDL cholesterol reduction with statins: the evidence, etiology, and therapeutic challenges. Curr Atheroscler Rep 14(1):1–10.

[3] Ridker, P.M., 2016. Residual inflammatory risk: addressing the obverse side of the atherosclerosis prevention coin. Eur Heart J 37(22):1720–2.

[4] Fisher, E.A., J.E. Feig, B. Hewing, S.L. Hazen, and J.D. Smith, 2012. High-Density Lipoprotein Function, Dysfunction, and Reverse Cholesterol Transport. Arteriosclerosis Thrombosis and Vascular Biology 32(12):2813–2820.

[5] Rohatgi, A., A. Khera, J.D. Berry, E.G. Givens, C.R. Ayers, K.E. Wedin, et al., 2014. HDL cholesterol efflux capacity and incident cardiovascular events. N Engl J Med 371(25):2383–93.

[6] Saleheen, D., R. Scott, S. Javad, W. Zhao, A. Rodrigues, A. Picataggi, et al., 2015. Association of HDL cholesterol efflux capacity with incident coronary heart disease events: a prospective case-control study. Lancet Diabetes Endocrinol 3(7):507–13.

[7] Rosenson, R.S., H.B. Brewer, Jr., B.J. Ansell, P. Barter, M.J. Chapman, J.W. Heinecke, et al., 2016. Dysfunctional HDL and atherosclerotic cardiovascular disease. Nat Rev Cardiol 13(1):48–60.

[8] Feig, J.E., B. Hewing, J.D. Smith, S.L. Hazen, and E.A. Fisher, 2014. High-density lipoprotein and atherosclerosis regression: evidence from preclinical and clinical studies. Circ Res 114(1):205–13.

[9] Huang, J., D. Wang, L.H. Huang, and H. Huang, 2020. Roles of Reconstituted High-Density Lipoprotein Nanoparticles in Cardiovascular Disease: A New Paradigm for Drug Discovery. Int J Mol Sci 21(3).

[10] Siebel, A.L., S.E. Heywood, and B.A. Kingwell, 2015. HDL and glucose metabolism: current evidence and therapeutic potential. Front Pharmacol 6:258.

[11] Dalla-Riva, J., K.G. Stenkula, J. Petrlova, and J.O. Lagerstedt, 2013. Discoidal HDL and apoA-I-derived peptides improve glucose uptake in skeletal muscle. Journal of Lipid Research 54(5):1275–82.

[12] Drew, B.G., S.J. Duffy, M.F. Formosa, A.K. Natoli, D.C. Henstridge, S.A. Penfold, et al., 2009. High-density lipoprotein modulates glucose metabolism in patients with type 2 diabetes mellitus. Circulation 119(15):2103–11.

[13] Shao, B.H., C.R. Tang, J.W. Heinecke, and J.F. Oram, 2010. Oxidation of apolipoprotein A-I by myeloperoxidase impairs the initial interactions with ABCA1 required for signaling and cholesterol export. Journal of Lipid Research 51(7):1849–1858.

[14] Huang, Y., J.A. DiDonato, B.S. Levison, D. Schmitt, L. Li, Y. Wu, et al., 2014. An abundant dysfunctional apolipoprotein A1 in human atheroma. Nat Med 20(2): 193–203.

[15] Shao, B., S. Pennathur, and J.W. Heinecke, 2012. Myeloperoxidase targets apolipoprotein A-I, the major high density lipoprotein protein, for site-specific oxidation in human atherosclerotic lesions. Journal of Biological Chemistry 287(9):6375–86.

[16] Gonen, A., L.F. Hansen, W.W. Turner, E.N. Montano, X. Que, A. Rafia, et al., 2014. Atheroprotective immunization with malondialdehyde-modified LDL is hapten specific and dependent on advanced MDA adducts: implications for development of an atheroprotective vaccine. Journal of Lipid Research 55(10):2137–55.

[17] Busch, C.J. and C.J. Binder, 2017. Malondialdehyde epitopes as mediators of sterile inflammation. Biochim Biophys Acta 1862(4):398–406.

[18] Shanmugam, N., J.L. Figarola, Y. Li, P.M. Swiderski, S. Rahbar, and R. Natarajan, 2008. Proinflammatory effects of advanced lipoxidation end products in monocytes. Diabetes 57(4):879–88.

[19] Huang, J., P.G. Yancey, H. Tao, M.S. Borja, L.E. Smith, V. Kon, et al., 2020. Reactive Dicarbonyl Scavenging Effectively Reduces MPO-Mediated Oxidation of HDL and Restores PON1 Activity. Nutrients 12(7).

[20] May-Zhang, L.S., V. Yermalitsky, J. Huang, T. Pleasent, M.S. Borja, M.N. Oda, et al., 2018. Modification by isolevuglandins, highly reactive gamma-ketoaldehydes, deleteriously alters high-density lipoprotein structure and function. Journal of Biological Chemistry 293(24):9176–9187.

[21] Shao, B., S. Pennathur, I. Pagani, M.N. Oda, J.L. Witztum, J.F. Oram, et al., 2010. Modifying apolipoprotein A-I by malondialdehyde, but not by an array of other reactive carbonyls, blocks cholesterol efflux by the ABCA1 pathway. Journal of Biological Chemistry 285(24):18473–84.

[22] May-Zhang, L.S., V. Yermalitsky, J.T. Melchior, J. Morris, K.A. Tallman, M.S. Borja, et al., 2019. Modified sites and functional consequences of 4-oxo-2-nonenal adducts in HDL that are elevated in familial hypercholesterolemia. Journal of Biological Chemistry 294(50):19022–19033.

[23] Besler, C., K. Heinrich, L. Rohrer, C. Doerries, M. Riwanto, D.M. Shih, et al., 2011. Mechanisms underlying adverse effects of HDL on eNOS-activating pathways in patients with coronary artery disease. J Clin Invest 121(7):2693–708.

[24] Davies, S.S., E.J. Brantley, P.A. Voziyan, V. Amarnath, I. Zagol-Ikapitte, O. Boutaud, et al., 2006. Pyridoxamine analogues scavenge lipid-derived gamma-ketoaldehydes and protect against H2O2-mediated cytotoxicity. Biochemistry 45(51):15756–67.

[25] Huang, J., F. Smith, and P. Panizzi, 2014. Ordered cleavage of myeloperoxidase ester bonds releases active site heme leading to inactivation of myeloperoxidase by benzoic acid hydrazide analogs. Arch Biochem Biophys 548:74–85.

[26] Huang, J., F. Smith, J.R. Panizzi, D.C. Goodwin, and P. Panizzi, 2015. Inactivation of myeloperoxidase by benzoic acid hydrazide. Arch Biochem Biophys 570:14–22.

[27] Linton, M.F., J.B. Atkinson, and S. Fazio, 1995. Prevention of atherosclerosis in apolipoprotein E-deficient mice by bone marrow transplantation. Science 267(5200):1034–7.

[28] Nicholls, S.J., G. Ruotolo, H.B. Brewer, J.P. Kane, M.D. Wang, K.A. Krueger, et al., 2015. Cholesterol Efflux Capacity and Pre-Beta-1 HDL Concentrations Are Increased in Dyslipidemic Patients Treated With Evacetrapib. J Am Coll Cardiol 66(20):2201–2210.

[29] Khera, A.V., M. Cuchel, M. de la Llera-Moya, A. Rodrigues, M.F. Burke, K. Jafri, et al., 2011. Cholesterol efflux capacity, high-density lipoprotein function, and atherosclerosis. N Engl J Med 364(2):127–35.

[30] Hung, A.M., Y. Tsuchida, K.L. Nowak, S. Sarkar, M. Chonchol, V. Whitfield, et al., 2019. IL-1 Inhibition and Function of the HDL-Containing Fraction of Plasma in Patients with Stages 3 to 5 CKD. Clin J Am Soc Nephrol 14(5):702–711.

[31] Robinet, P., Z. Wang, S.L. Hazen, and J.D. Smith, 2010. A simple and sensitive enzymatic method for cholesterol quantification in macrophages and foam cells. Journal of Lipid Research 51(11):3364–9.

[32] Shea, S., J.H. Stein, N.W. Jorgensen, R.L. McClelland, L. Tascau, S. Shrager, et al., 2019. Cholesterol Mass Efflux Capacity, Incident Cardiovascular Disease, and Progression of Carotid Plaque. Arterioscler Thromb Vasc Biol 39(1):89–96.

[33] Furuhashi, M., G. Tuncman, C.Z. Gorgun, L. Makowski, G. Atsumi, E. Vaillancourt, et al., 2007. Treatment of diabetes and atherosclerosis by inhibiting fatty-acid-binding protein aP2. Nature 447(7147):959–65.

[34] Liang, W., A.L. Menke, A. Driessen, G.H. Koek, J.H. Lindeman, R. Stoop, et al., 2014. Establishment of a general NAFLD scoring system for rodent models and comparison to human liver pathology. PLoS One 9(12):e115922.

[35] Daugherty, A., A.R. Tall, M. Daemen, E. Falk, E.A. Fisher, G. Garcia-Cardena, et al., 2017. Recommendation on Design, Execution, and Reporting of Animal Atherosclerosis Studies: A Scientific Statement From the American Heart Association. Arterioscler Thromb Vasc Biol 37(9):e131–e157.

[36] Tao, H., J. Huang, P.G. Yancey, V. Yermalitsky, J.L. Blakemore, Y. Zhang, et al., 2020. Scavenging of reactive dicarbonyls with 2-hydroxybenzylamine reduces atherosclerosis in hypercholesterolemic Ldlr(-/-) mice. Nature Communications 11(1):4084.

[37] Babaev, V.R., L. Ding, Y. Zhang, J.M. May, S.A. Ramsey, K.C. Vickers, et al., 2018. Loss of 2 Akt (Protein Kinase B) Isoforms in Hematopoietic Cells Diminished Monocyte and Macrophage Survival and Reduces Atherosclerosis in Ldl Receptor-Null Mice. Arterioscler Thromb Vasc Biol:ATVBAHA118312206.

[38] Maessen, D.E., O. Brouwers, K.H. Gaens, K. Wouters, J.P. Cleutjens, B.J. Janssen, et al., 2016. Delayed Intervention With Pyridoxamine Improves Metabolic Function and Prevents Adipose Tissue Inflammation and Insulin Resistance in High-Fat Diet-Induced Obese Mice. Diabetes 65(4):956–66.

[39] Tao, H., P.G. Yancey, V.R. Babaev, J.L. Blakemore, Y. Zhang, L. Ding, et al., 2015. Macrophage SR-BI mediates efferocytosis via Src/PI3K/Rac1 signaling and reduces atherosclerotic lesion necrosis. Journal of Lipid Research 56(8):1449–60.

[40] Su, Y.R., M.F. Linton, and S. Fazio, 2002. Rapid quantification of murine ABC mRNAs by real time reverse transcriptase-polymerase chain reaction. J Lipid Res 43(12):2180–7.

[41] Madu, H., J. Avance, S. Chetyrkin, C. Darris, K.L. Rose, O.A. Sanchez, et al., 2015. Pyridoxamine protects proteins from damage by hypohalous acids in vitro and in vivo. Free Radic Biol Med 89:83–90.

[42] Yla-Herttuala, S., W. Palinski, M. Rosenfeld, S. Parthasarathy, T. Carew, S. Butler, et al., 1989. Evidence for the presence of oxidatively modified low density lipoprotein in atherosclerotic lesions of rabbit and man. J Clin Invest 84:1086–1095.

[43] Shaw, P.X., S. Horkko, S. Tsimikas, M.K. Chang, W. Palinski, G.J. Silverman, et al., 2001. Human-derived anti-oxidized LDL autoantibody blocks uptake of oxidized LDL by macrophages and localizes to atherosclerotic lesions in vivo. Arterioscler Thromb Vasc Biol 21(8):1333–9.

[44] Schiopu, A., B. Frendeus, B. Jansson, I. Soderberg, I. Ljungcrantz, Z. Araya, et al., 2007. Recombinant antibodies to an oxidized low-density lipoprotein epitope induce rapid regression of atherosclerosis in apobec-1(-/-)/low-density lipoprotein receptor(-/-) mice. J Am Coll Cardiol 50(24):2313–8.

[45] Merat, S., F. Casanada, M. Sutphin, W. Palinski, and P.D. Reaven, 1999. Western-type diets induce insulin resistance and hyperinsulinemia in LDL receptor-deficient mice but do not increase aortic atherosclerosis compared with normoinsulinemic mice in which similar plasma cholesterol levels are achieved by a fructose-rich diet. Arterioscler Thromb Vasc Biol 19(5):1223–30.

[46] Hartvigsen, K., C.J. Binder, L.F. Hansen, A. Rafia, J. Juliano, S. Horkko, et al., 2007. A diet-induced hypercholesterolemic murine model to study atherogenesis without obesity and metabolic syndrome. Arterioscler Thromb Vasc Biol 27(4):878–85.

[47] Li, A.C., K.K. Brown, M.J. Silvestre, T.M. Willson, W. Palinski, and C.K. Glass, 2000. Peroxisome proliferator-activated receptor gamma ligands inhibit development of atherosclerosis in LDL receptor-deficient mice. Journal of Clinical Investigation 106(4):523–531.

[48] Arnold, A.P., L.A. Cassis, M. Eghbali, K. Reue, and K. Sandberg, 2017. Sex Hormones and Sex Chromosomes Cause Sex Differences in the Development of Cardiovascular Diseases. Arterioscler Thromb Vasc Biol 37(5):746–756.

[49] Warnatsch, A., M. Ioannou, Q. Wang, and V. Papayannopoulos, 2015. Inflammation. Neutrophil extracellular traps license macrophages for cytokine production in atherosclerosis. Science 349(6245):316–20.

[50] Odobasic, D., A.R. Kitching, and S.R. Holdsworth, 2016. Neutrophil-Mediated Regulation of Innate and Adaptive Immunity: The Role of Myeloperoxidase. J Immunol Res 2016:2349817.

[51] Swirski, F.K., P. Libby, E. Aikawa, P. Alcaide, F.W. Luscinskas, R. Weissleder, et al., 2007. Ly-6Chi monocytes dominate hypercholesterolemia-associated monocytosis and give rise to macrophages in atheromata. J Clin Invest 117(1):195–205.

[52] Babaev, V.R., K.E. Hebron, C.B. Wiese, C.L. Toth, L. Ding, Y. Zhang, et al., 2014. Macrophage deficiency of Akt2 reduces atherosclerosis in Ldlr null mice. Journal of Lipid Research 55(11):2296–308.

[53] Gao, M., D. Zhao, S. Schouteden, M.G. Sorci-Thomas, P.P. Van Veldhoven, K. Eggermont, et al., 2014. Regulation of high-density lipoprotein on hematopoietic stem/progenitor cells in atherosclerosis requires scavenger receptor type BI expression. Arterioscler Thromb Vasc Biol 34(9):1900–9.

[54] Roberts, L.J., J.A. Oates, M.F. Linton, S. Fazio, B.P. Meador, M.D. Gross, et al., 2007. The relationship between dose of vitamin E and suppression of oxidative stress in humans. Free Radical Biology and Medicine 43(10):1388–1393.

[55] May-Zhang, L.S., A. Kirabo, J. Huang, M.F. Linton, S.S. Davies, and K.T. Murray, 2020. Scavenging Reactive Lipids to Prevent Oxidative Injury. Annu Rev Pharmacol Toxicol.

[56] Delporte, C., P. Van Antwerpen, L. Vanhamme, T. Roumeguere, and K. Zouaoui Boudjeltia, 2013. Low-density lipoprotein modified by myeloperoxidase in inflammatory pathways and clinical studies. Mediators Inflamm 2013:971579.

[57] Panzenboeck, U., S. Raitmayer, H. Reicher, H. Lindner, O. Glatter, E. Malle, et al., 1997. Effects of reagent and enzymatically generated hypochlorite on physicochemical and metabolic properties of high density lipoproteins. Journal of Biological Chemistry 272(47):29711–20.

[58] Takata, K., S. Horiuchi, and Y. Morino, 1989. Scavenger receptor-mediated recognition of maleylated albumin and its relation to subsequent endocytic degradation. Biochim Biophys Acta 984(3):273–80.

[59] Kimball, A., M. Schaller, A. Joshi, F.M. Davis, A. denDekker, A. Boniakowski, et al., 2018. Ly6C(Hi) Blood Monocyte/Macrophage Drive Chronic Inflammation and Impair Wound Healing in Diabetes Mellitus. Arterioscler Thromb Vasc Biol 38(5):1102–1114.

[60] Berg, K.E., I. Ljungcrantz, L. Andersson, C. Bryngelsson, B. Hedblad, G.N. Fredrikson, et al., 2012. Elevated CD14++CD16-monocytes predict cardiovascular events. Circ Cardiovasc Genet 5(1):122–31.

[61] Heine, G.H., C. Ulrich, E. Seibert, S. Seiler, J. Marell, B. Reichart, et al., 2008. CD14(++)CD16+ monocytes but not total monocyte numbers predict cardiovascular events in dialysis patients. Kidney Int 73(5):622–9.

[62] Fadini, G.P., F. Simoni, R. Cappellari, N. Vitturi, S. Galasso, S. Vigili de Kreutzenberg, et al., 2014. Pro-inflammatory monocyte-macrophage polarization imbalance in human hypercholesterolemia and atherosclerosis. Atherosclerosis 237(2):805–8.

[63] Hjuler Nielsen, M., H. Irvine, S. Vedel, B. Raungaard, H. Beck-Nielsen, and A. Handberg, 2015. Elevated atherosclerosis-related gene expression, monocyte activation and microparticle-release are related to increased lipoprotein-associated oxidative stress in familial hypercholesterolemia. PLoS One 10(4):e0121516.

[64] Rogacev, K.S., B. Cremers, A.M. Zawada, S. Seiler, N. Binder, P. Ege, et al., 2012. CD14++CD16+ monocytes independently predict cardiovascular events: a cohort study of 951 patients referred for elective coronary angiography. J Am Coll Cardiol 60(16):1512–20.

[65] Tie, G., K.E. Messina, J. Yan, J.A. Messina, and L.M. Messina, 2014. Hypercholesterolemia induces oxidant stress that accelerates the ageing of hematopoietic stem cells. J Am Heart Assoc 3(1):e000241.

[66] van der Valk, F.M., C. Kuijk, S.L. Verweij, L.C.A. Stiekema, Y. Kaiser, S. Zeerleder, et al., 2017. Increased haematopoietic activity in patients with atherosclerosis. Eur Heart J 38(6):425–432.

[67] Schill, R.L., D.A. Knaack, H.R. Powers, Y. Chen, M. Yang, D.J. Schill, et al., 2020. Modification of HDL by reactive aldehydes alters select cardioprotective functions of HDL in macrophages. FEBS J 287(4):695–707.

[68] Tall, A.R., L. Yvan-Charvet, M. Westerterp, and A.J. Murphy, 2012. Cholesterol efflux: a novel regulator of myelopoiesis and atherogenesis. Arterioscler Thromb Vasc Biol 32(11):2547–52.

[69] Nahrendorf, M. and F.K. Swirski, 2016. Abandoning M1/M2 for a Network Model of Macrophage Function. Circ Res 119(3):414–7.

[70] Ridker, P.M., B.M. Everett, T. Thuren, J.G. MacFadyen, W.H. Chang, C. Ballantyne, et al., 2017. Antiinflammatory Therapy with Canakinumab for Atherosclerotic Disease. N Engl J Med 377(12):1119–1131.

[71] Boring, L., J. Gosling, M. Cleary, and I.F. Charo, 1998. Decreased lesion formation in CCR2-/-mice reveals a role for chemokines in the initiation of atherosclerosis. Nature 394(6696):894–7.

[72] Bartlett, B., H.P. Ludewick, A. Misra, S. Lee, and G. Dwivedi, 2019. Macrophages and T cells in atherosclerosis: a translational perspective. Am J Physiol Heart Circ Physiol 317(2):H375–H386.

[73] Doran, A.C., 2022. Inflammation Resolution: Implications for Atherosclerosis. Circ Res 130(1):130–148.

[74] Yurdagul, A., Jr., M. Subramanian, X. Wang, S.B. Crown, O.R. Ilkayeva, L. Darville, et al., 2020. Macrophage Metabolism of Apoptotic Cell-Derived Arginine Promotes Continual Efferocytosis and Resolution of Injury. Cell Metab 31(3):518-533 e10.

[75] Papac-Milicevic, N., C.J. Busch, and C.J. Binder, 2016. Malondialdehyde Epitopes as Targets of Immunity and the Implications for Atherosclerosis. Adv Immunol 131:1–59.

[76] Lee, M.K., X.L. Moore, Y. Fu, A. Al-Sharea, D. Dragoljevic, M.A. Fernandez-Rojo, et al., 2016. High-density lipoprotein inhibits human M1 macrophage polarization through redistribution of caveolin-1. Br J Pharmacol 173(4):741–51.

[77] Sanson, M., E. Distel, and E.A. Fisher, 2013. HDL induces the expression of the M2 macrophage markers arginase 1 and Fizz-1 in a STAT6-dependent process. PLoS One 8(8):e74676.

[78] Yvan-Charvet, L., C. Welch, T.A. Pagler, M. Ranalletta, M. Lamkanfi, S. Han, et al., 2008. Increased inflammatory gene expression in ABC transporter-deficient macrophages: free cholesterol accumulation, increased signaling via toll-like receptors, and neutrophil infiltration of atherosclerotic lesions. Circulation 118(18):1837–47.

[79] Zhu, X., J.Y. Lee, J.M. Timmins, J.M. Brown, E. Boudyguina, A. Mulya, et al., 2008. Increased cellular free cholesterol in macrophage-specific Abca1 knock-out mice enhances pro-inflammatory response of macrophages. Journal of Biological Chemistry 283(34):22930–41.

[80] Groenen, A.G., B. Halmos, A.R. Tall, and M. Westerterp, 2021. Cholesterol efflux pathways, inflammation, and atherosclerosis. Crit Rev Biochem Mol Biol 56(4):426–439.

[81] Henriksen, E.J., M.K. Diamond-Stanic, and E.M. Marchionne, 2011. Oxidative stress and the etiology of insulin resistance and type 2 diabetes. Free Radic Biol Med 51(5):993–9.

[82] Vidal, N., J.P. Cavaille, M. Poggi, F. Peiretti, and P. Stocker, 2012. A nonradioisotope chemiluminescent assay for evaluation of 2-deoxyglucose uptake in 3T3-L1 adipocytes. Effect of various carbonyls species on insulin action. Biochimie 94(12):2569–76.

[83] Pillon, N.J., M.L. Croze, R.E. Vella, L. Soulere, M. Lagarde, and C.O. Soulage, 2012. The lipid peroxidation by-product 4-hydroxy-2-nonenal (4-HNE) induces insulin resistance in skeletal muscle through both carbonyl and oxidative stress. Endocrinology 153(5):2099–111.

[84] Shearn, C.T., K.S. Fritz, P. Reigan, and D.R. Petersen, 2011. Modification of Akt2 by 4-hydroxynonenal inhibits insulin-dependent Akt signaling in HepG2 cells. Biochemistry 50(19):3984–96.

[85] de Haan, W., A. Bhattacharjee, P. Ruddle, M.H. Kang, and M.R. Hayden, 2014. ABCA1 in adipocytes regulates adipose tissue lipid content, glucose tolerance, and insulin sensitivity. Journal of Lipid Research 55(3):516–23.

[86] Sanchez-Aguilera, P., A. Diaz-Vegas, C. Campos, O. Quinteros-Waltemath, H. Cerda-Kohler, G. Barrientos, et al., 2018. Role of ABCA1 on membrane cholesterol content, insulin-dependent Akt phosphorylation and glucose uptake in adult skeletal muscle fibers from mice. Biochim Biophys Acta Mol Cell Biol Lipids 1863(12):1469–1477.

[87] Edmunds, S.J., R. Liebana-Garcia, O. Nilsson, J. Domingo-Espin, C. Gronberg, K.G. Stenkula, et al., 2019. ApoAI-derived peptide increases glucose tolerance and prevents formation of atherosclerosis in mice. Diabetologia 62(7):1257–1267.

[88] Abdullah, K.M., F. Abul Qais, H. Hasan, and I. Naseem, 2019. Anti-diabetic study of vitamin B6 on hyperglycaemia induced protein carbonylation, DNA damage and ROS production in alloxan induced diabetic rats. Toxicol Res (Camb) 8(4):568–579.

