## Supplemental Material for "Scavenging dicarbonyls with 5’-O-pentyl-pyridoxamine increases HDL net cholesterol efflux capacity and attenuates atherosclerosis and insulin resistance"

Short title: 5'-O-pentyl-pyridoxamine improves insulin sensitivity and reduces atherosclerosis

#### Figure S1

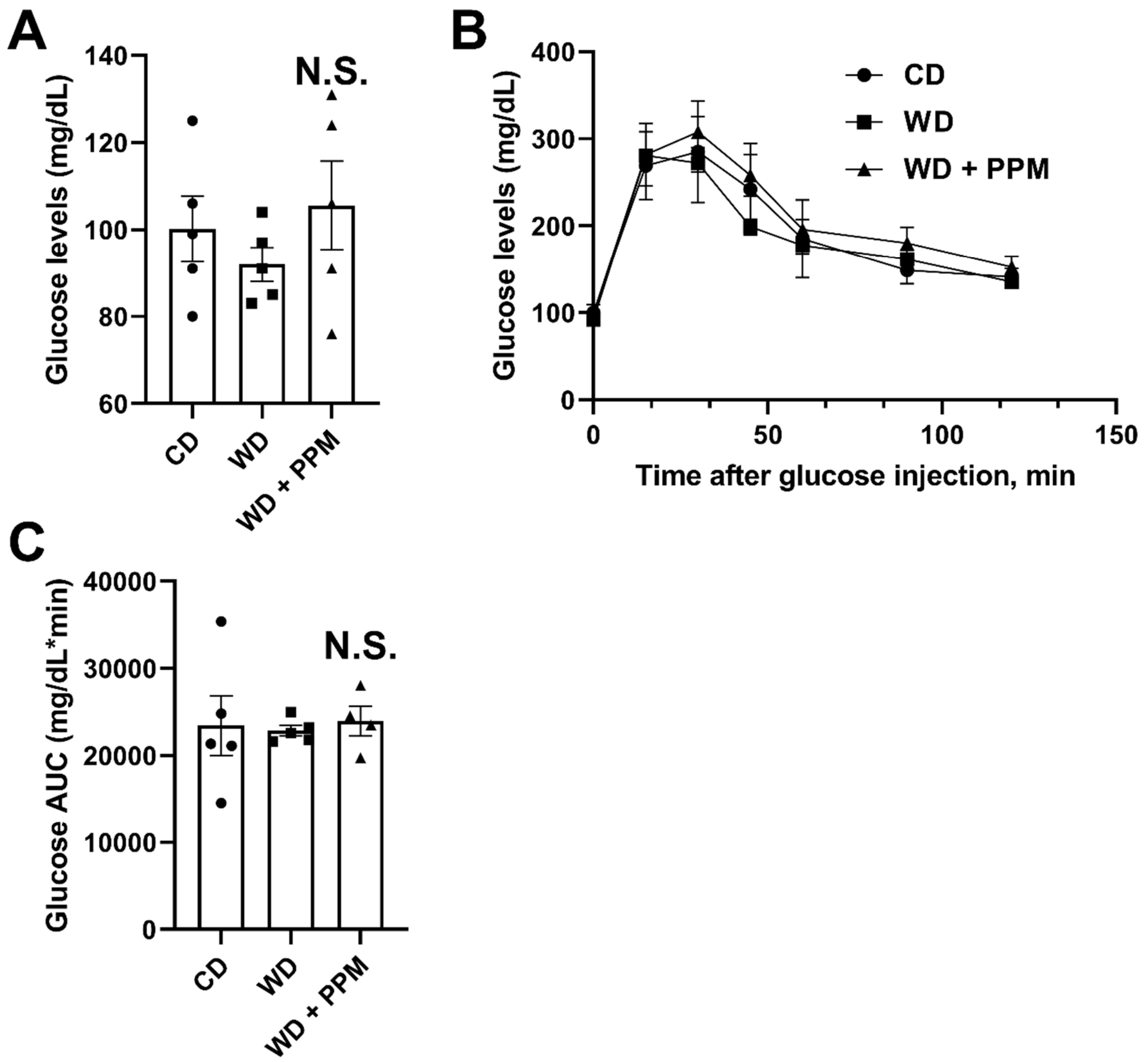


**Figure S1. PPM does not impact fasting glucose levels or response to glucose challenge in female *Ldlr*$^{-/-}$ mice.** A. PPM has no effect on fasting glucose levels in female *Ldlr*$^{-/-}$ mice. Fasting glucose was measured in female *Ldlr*$^{-/-}$ mice on a chow diet or on a western diet with and without PPM treatment for 16 weeks. B-C. Effect of PPM on glucose levels in female *Ldlr*$^{-/-}$ mice in response to injection of glucose using IPGTT as described in Methods (n=5 in each group). For each experiment, graph data are expressed as mean ± SEM; N.S. by one-way ANOVA with Bonferroni's post hoc test.

#### Figure S2

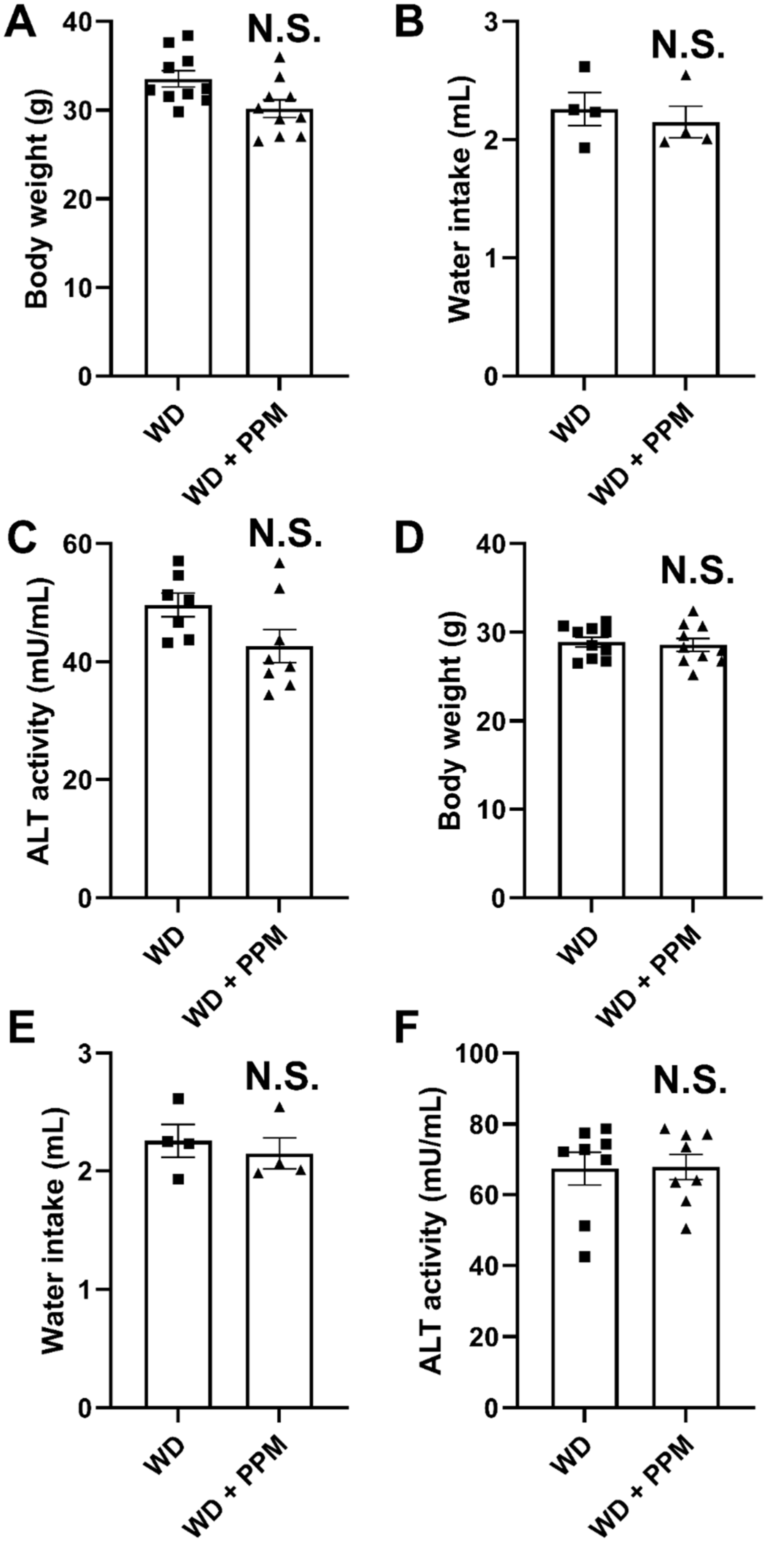


**Figure S2. PPM does not affect the body weight, water intake or plasma ALT activities in male or female *Ldlr*$^{-/-}$ mice.** A-F. The effect of PPM on the body weight, water intake and ALT activities in male (A-C) or female (D-F) *Ldlr*$^{-/-}$ mice on a western diet for 16 weeks. For each experiment, graph data are expressed as mean ± SEM; *P <0.05 by one-way ANOVA.

#### Figure S3

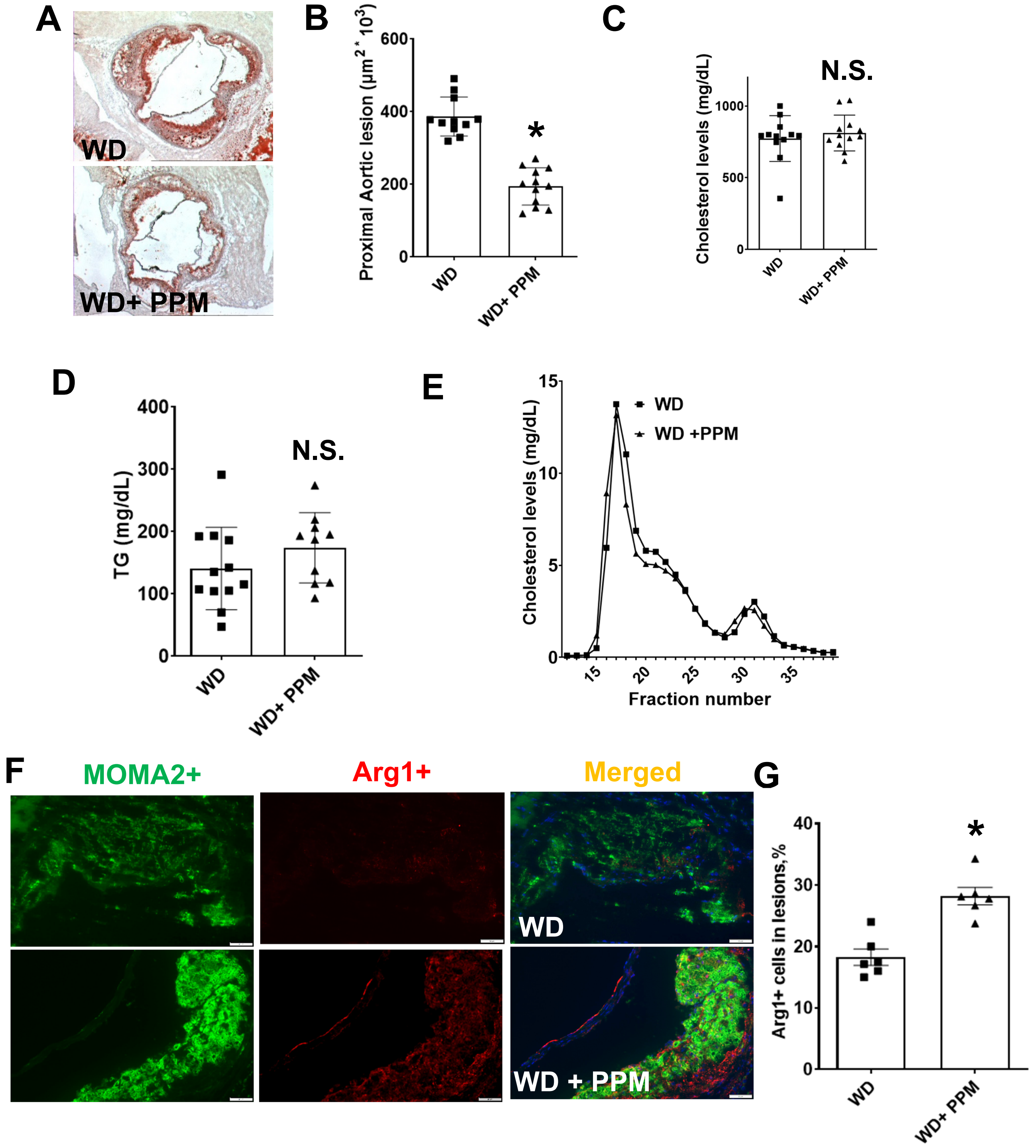


**Figure S3. PPM reduces atherosclerotic lesion extent and plaque inflammatory macrophage content in female *Ldlr*$^{-/-}$ mice**. A-C. PPM does not affect plasma cholesterol, triglycerides, or FPLC lipoprotein profile in female *Ldlr*$^{-/-}$ mice. D&E. PPM reduces the aortic root atherosclerotic lesions in female *Ldlr*$^{-/-}$ mice. Female *Ldlr*$^{-/-}$ mice were pretreated with vehicle alone or with 1 g/L of PPM for 2 weeks on chow diet. Then, mice were treated with vehicle or 1 g/L of PPM for 16 weeks on WD. Oil-Red-O staining of atherosclerotic lesions in female *Ldlr*$^{-/-}$ mice with vehicle (n=12) or PPM (n=12). Scale bar = 200 μm. F&G. Immunohistochemistry staining of Arg1+ macrophages was performed as described in Methods. G. PPM increases the number of Arg1+ macrophages in lesions of *Ldlr*$^{-/-}$ mice. F. Scale bar = 40 μm. n=6 in each group, for each experiment, graph data are expressed as mean ± SEM; *P <0.05 by two-sided unpaired t-test.

**Figure S4**

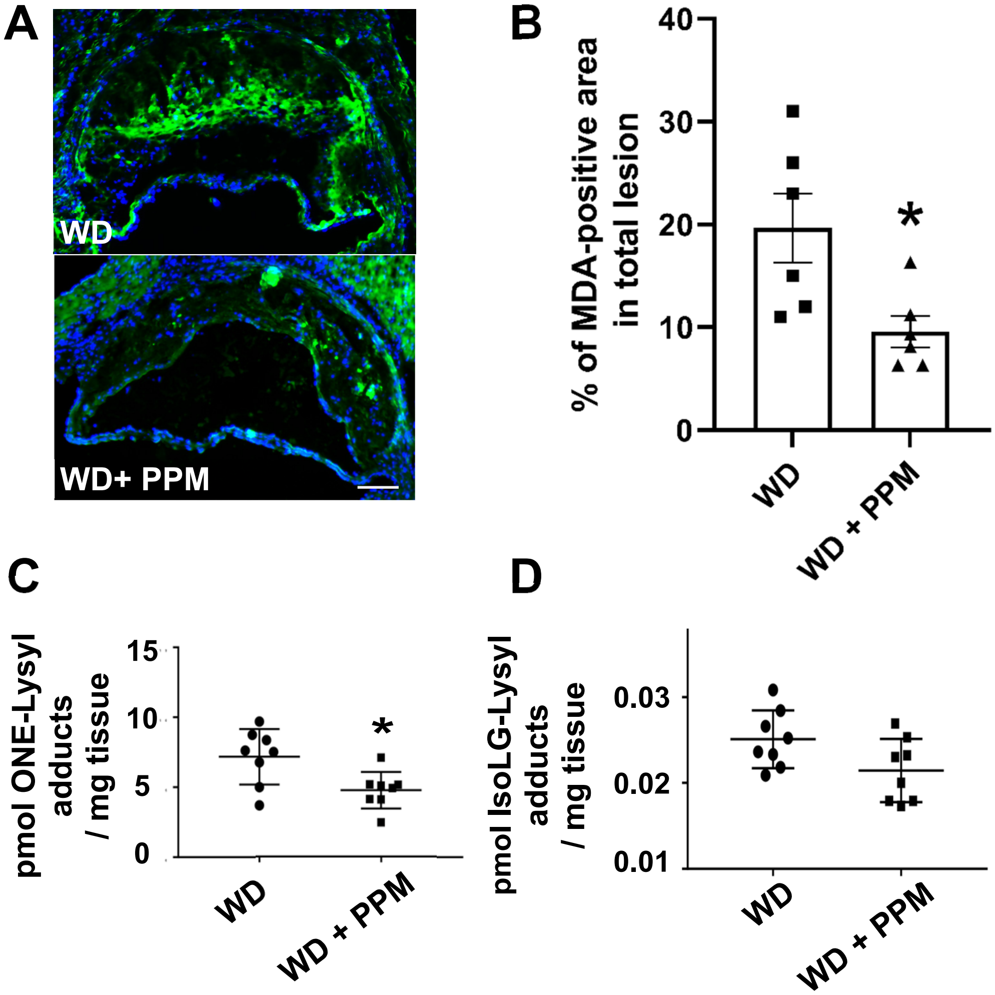


**Figure S4. PPM reduces the lipid reactive dicarbonyl content in lesions of female and male *Ldlr*$^{-/-}$ mice.** A-B. PPM reduces MDA-Lysyl adducts in the proximal aortic lesions of female *Ldlr*$^{-/-}$ mice. MDA adducts were measured by immunofluorescence using anti-MDA adduct primary antibody (green). Nuclei were counterstained with Hoechst (blue). Representative images (A) and quantitation (B) of MDA adduct /staining in proximal aortic root sections. N = 6 mice per group. Scale bar = 50 μm. n=6 in each group, for each experiment, graph data are expressed as mean ± SEM; *P <0.05 by two-sided unpaired t-test. C. PPM reduces the levels of ONE-Lysyl adducts in the aorta of male *Ldlr*$^{-/-}$ mice. Aortic tissues were isolated from male *Ldlr*$^{-/-}$ mice and ONE-Lysyl adducts were measured by LC/MS/MS. D. The effect of PPM on the levels of IsoLG-Lysyl adducts in the aorta of *Ldlr*$^{-/-}$ mice. Aortic tissues were isolated from the *Ldlr*$^{-/-}$ mice and IsoLG-Lysyl adducts were measured by LC/MS/MS. C-D. n=8 in each group, for each experiment, graph data are expressed as mean ± SEM; *P <0.05 by two-sided unpaired t-test.

### Figure S5

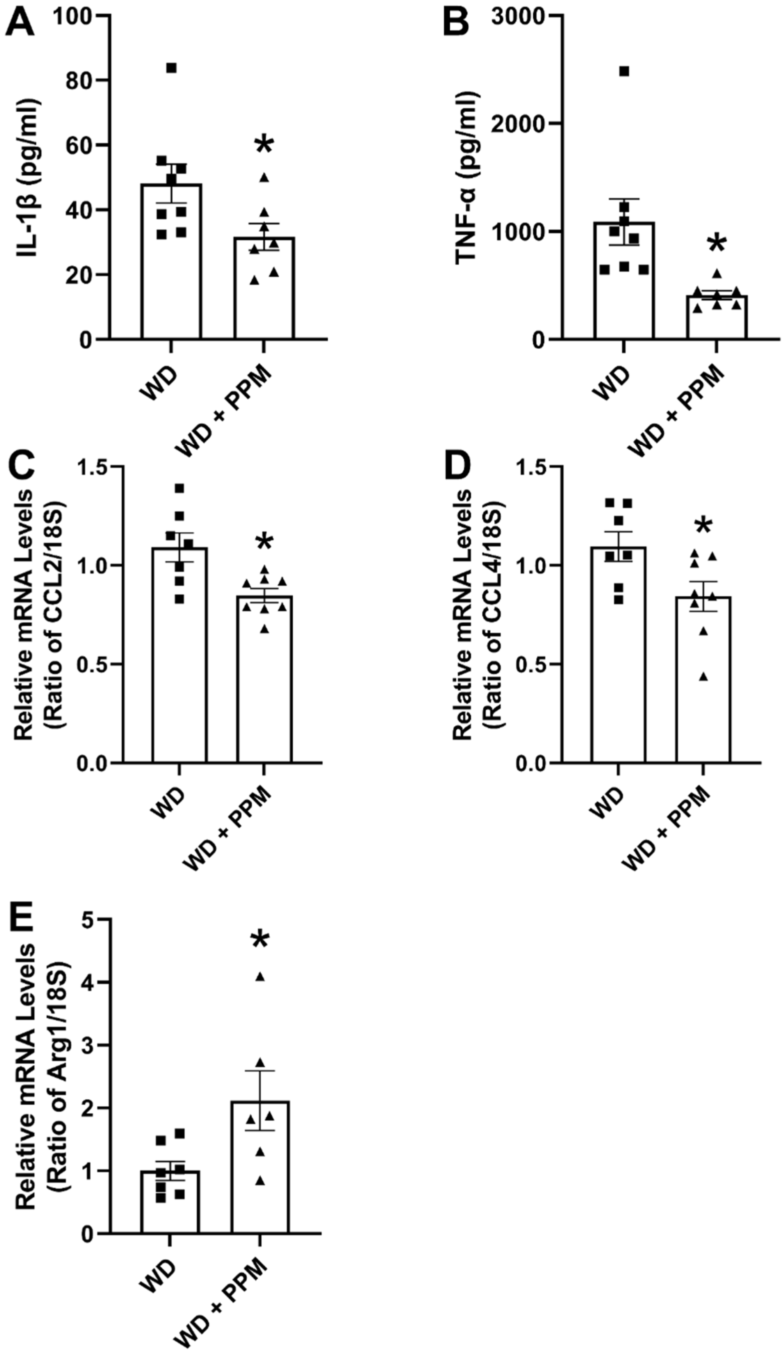


**Figure S5. PPM reduces serum and hepatic inflammatory markers in male *Ldlr$^{-/-}$* mice.** Male *Ldlr$^{-/-}$* mice were treated with 1 g/L of PPM for 16 weeks on WD. A-B. PPM reduces the serum levels of IL-1β and TNF-α, which were measured by ELISA (n=7 per group). C-E. PPM decreases the hepatic expression of inflammatory CCL2 and CCL4 and increases anti-inflammatory Arg1 mRNA levels (n=6, 7, or 8 per group). The mRNA levels were measured by real-time PCR as described in Methods. Data are expressed as mean ± SEM; *P <0.05 by two-sided unpaired t-test.

#### Figure S6

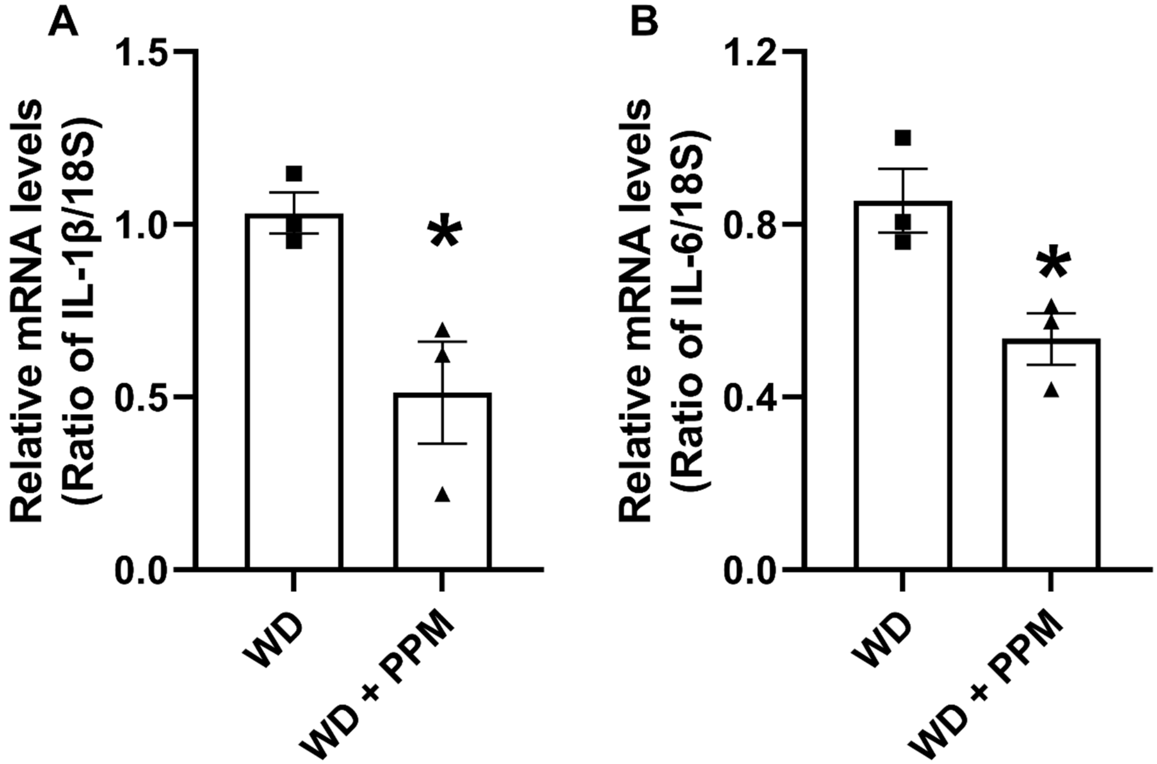


**Figure S6. PPM reduces the expression of IL-1β and IL-6 in lesions of male *Ldlr*$^{-/-}$ mice.** A-B. PPM decreases the mRNA levels of IL-1β and IL-6 in plaques of male *Ldlr*$^{-/-}$ mice. Aortic tissues were isolated from male *Ldlr*$^{-/-}$ mice that had been fed a western diet for 16 weeks, and the mRNA levels of IL-1β (A) and IL-6 (B) were measured by real-time PCR as described in Methods. n=3 per group. Data are expressed as mean ± SEM; *P <0.05 by two-sided unpaired t-test.

#### Figure S7

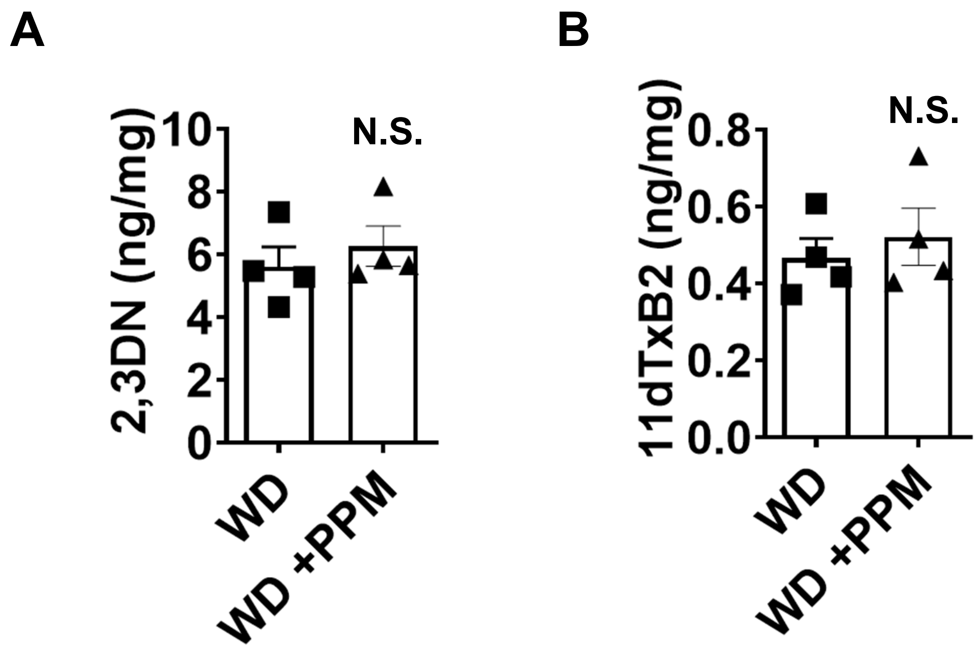


**Figure S7. PPM does not change the levels of PG metabolites in *Ldlr*$^{-/-}$ mice.** A-B. Male *Ldlr*$^{-/-}$ mice were given vehicle (n=8) or 1 mg/mL PPM (n=8) for 8 weeks. Urine was collected for 6 h in metabolic cages with 2 mice per cage after 8 weeks of treatment with PPM and analyzed by GC/MS for 2,3-dinor-6-ketoPGF1 (2,3DN) and 11-dehydro TxB2 (11dTxB2) by the Eicosanoid Analysis Core at Vanderbilt University. The creatinine levels are measured using a kit (Enzo Life Sciences). The urinary metabolite levels in each sample were normalized using the urinary creatinine level of the sample and expressed as ng/mg creatinine. PPM does not change the levels of 2,3DN or TxB2 in *Ldlr*$^{-/-}$ mice (n=4 in each group). Graph data are expressed as mean ± SEM, there is no significant difference between groups determined by using two-sided unpaired t-test.

#### Figure S8

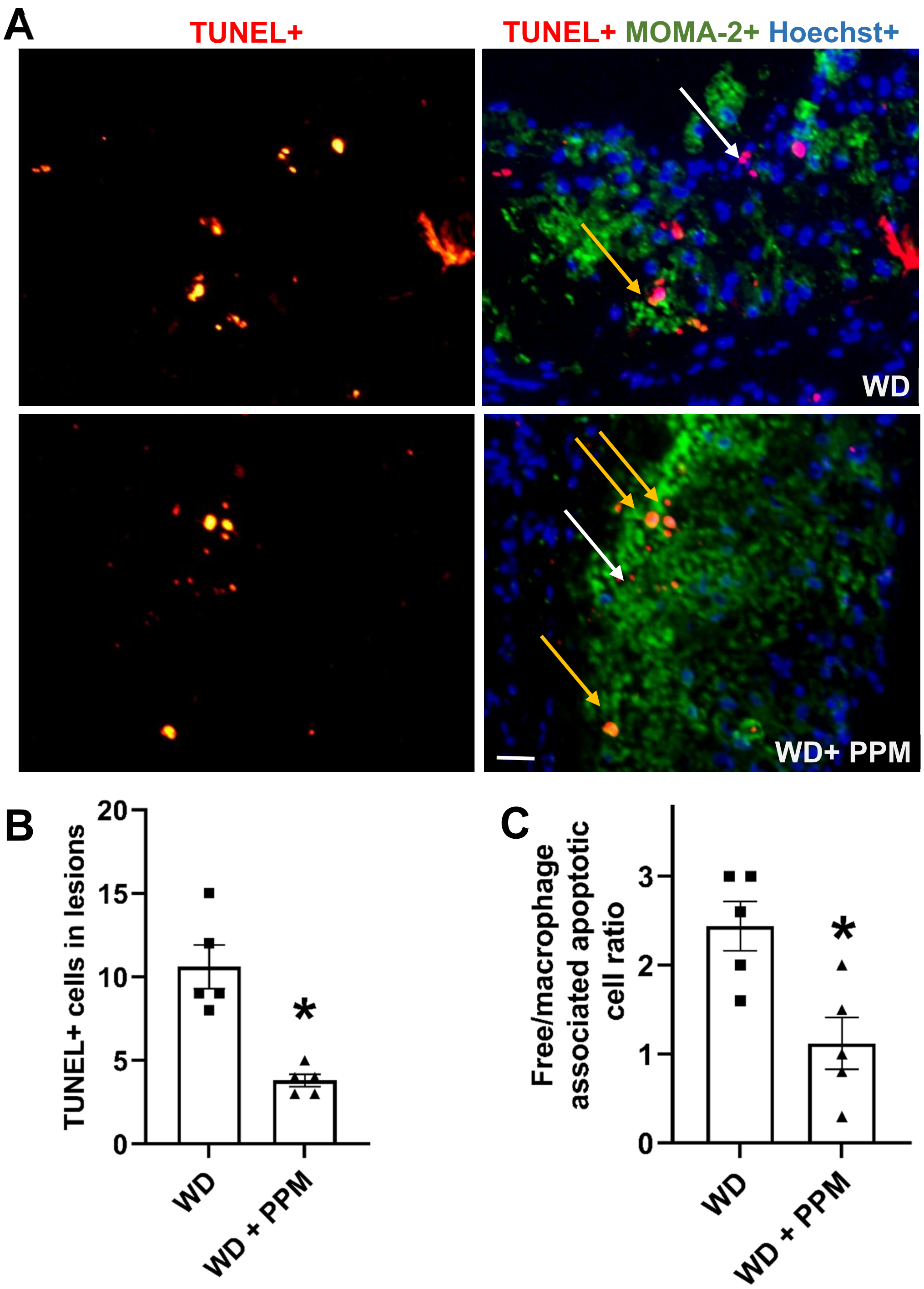


**Figure S8. PPM reduces the number of apoptotic cells and increases efferocytosis in atherosclerotic plaques of female *Ldlr*$^{-/-}$ mice.** A. Images depict TUNEL staining of nuclei (red) and merged images show TUNEL+, MOMA2+ (green) and Hoechst+ (blue) staining of atherosclerotic lesions in male *Ldlr*$^{-/-}$ mice. Yellow arrows show macrophage-associated TUNEL stain and white arrows depict free dead cells that were not associated with macrophages. Scale bar = 50 µm. B. Quantitation of the number of apoptotic cells. C. Efferocytosis was quantitated as the free vs macrophage-associated TUNEL-positive cells in the proximal aortic sections. B-C. n=6 per group. Graph data are expressed as mean ± SEM; *P <0.05 by two-sided unpaired T-test.

#### Figure S9

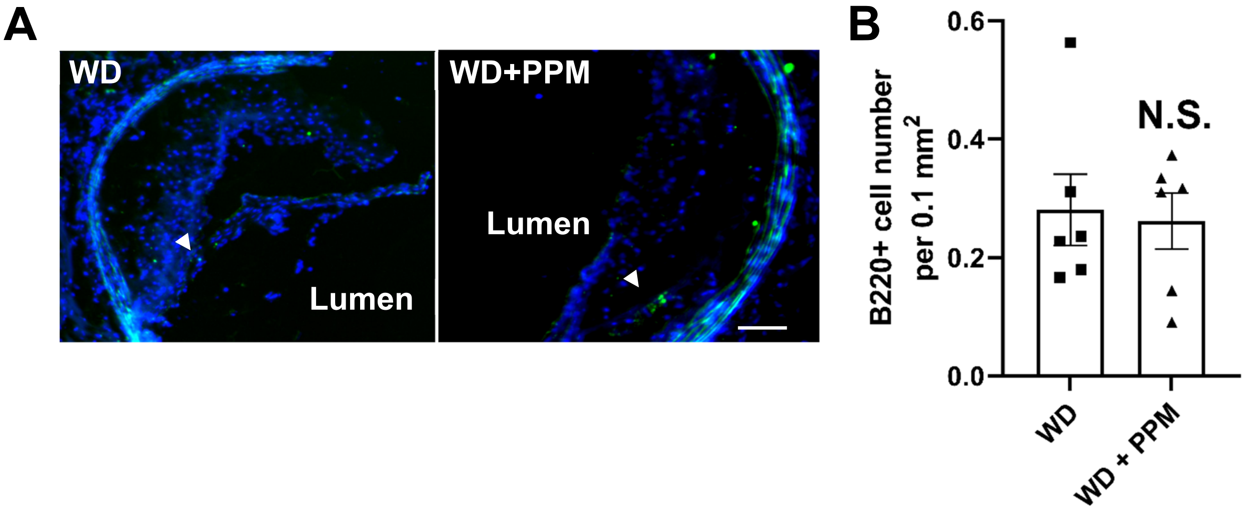


**Figure S9. PPM does not impact the number of B lymphocytes in plaques of female *Ldlr*$^{-/-}$ mice.** A-B. Immunohistochemistry staining and quantitation of B220+ (green) B lymphocytes in atherosclerotic lesions of female *Ldlr*$^{-/-}$ mice. Scale bar = 50 $\mu$m. n=6 or 7 per group. Graph data are expressed as mean $\pm$ SEM; N.S. by two-sided unpaired t-test.

### Figure S10

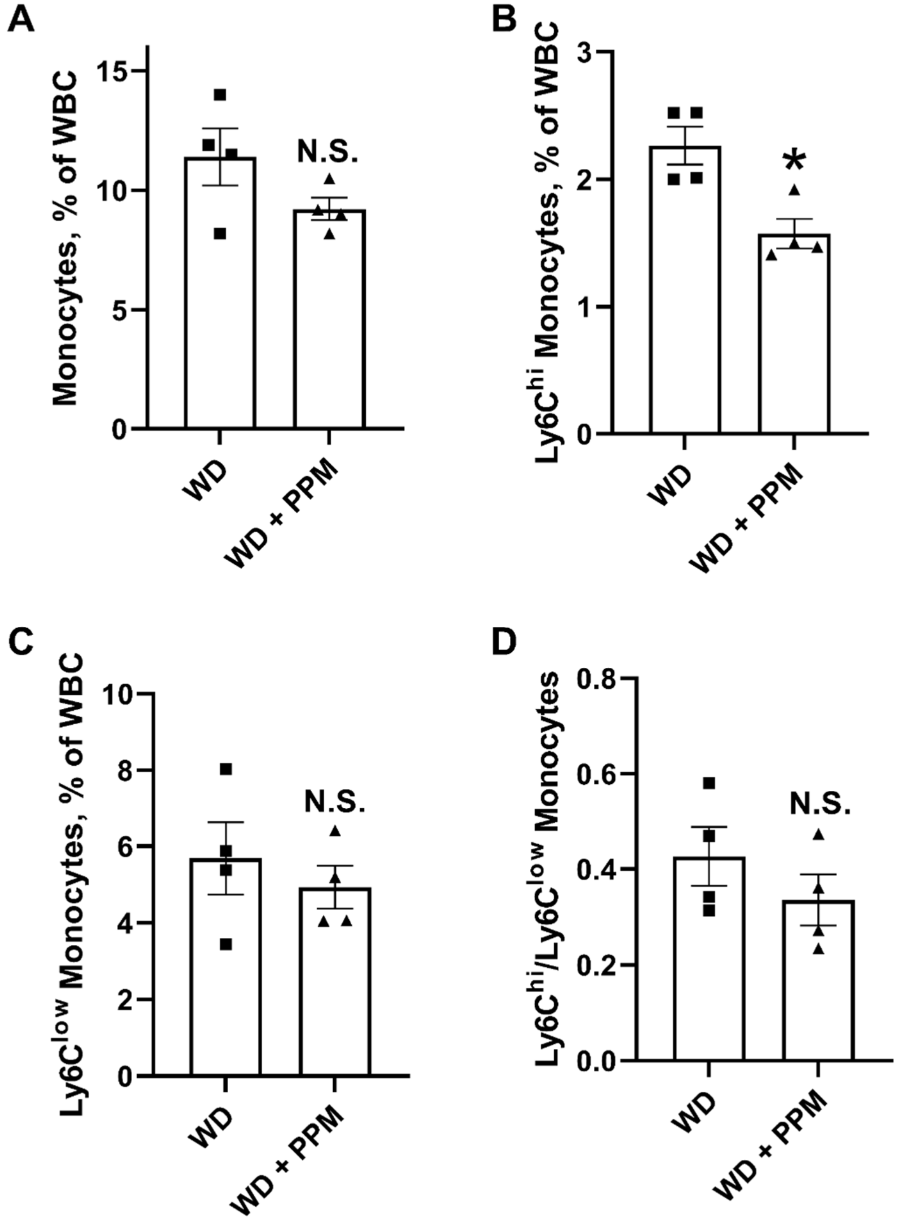


**Figure S10. PPM decreases the number of Ly6C$^{hi}$ monocytes in male *Ldlr$^{-/-}$* mice.** A-D. Male *Ldlr$^{-/-}$* mice were fed a western diet for 16 weeks and treated with water alone or with PPM. A. There was a trend toward a reduction in total blood monocytes in PPM versus water treated *Ldlr$^{-/-}$* mice. B. PPM reduced the number of Ly6C$^{hi}$ monocytes C. PPM did not impact the number of Ly6C $^{low}$ monocytes. F. There was a trend toward a reduction in the ratio of Ly6C$^{hi}$ to Ly6C $^{low}$ monocytes in PPM versus water treated *Ldlr$^{-/-}$* mice. n=8 in each group. Graph data are expressed as mean ± SEM; *P <0.05, N.S. by two-sided unpaired t-test.

### Figure S11

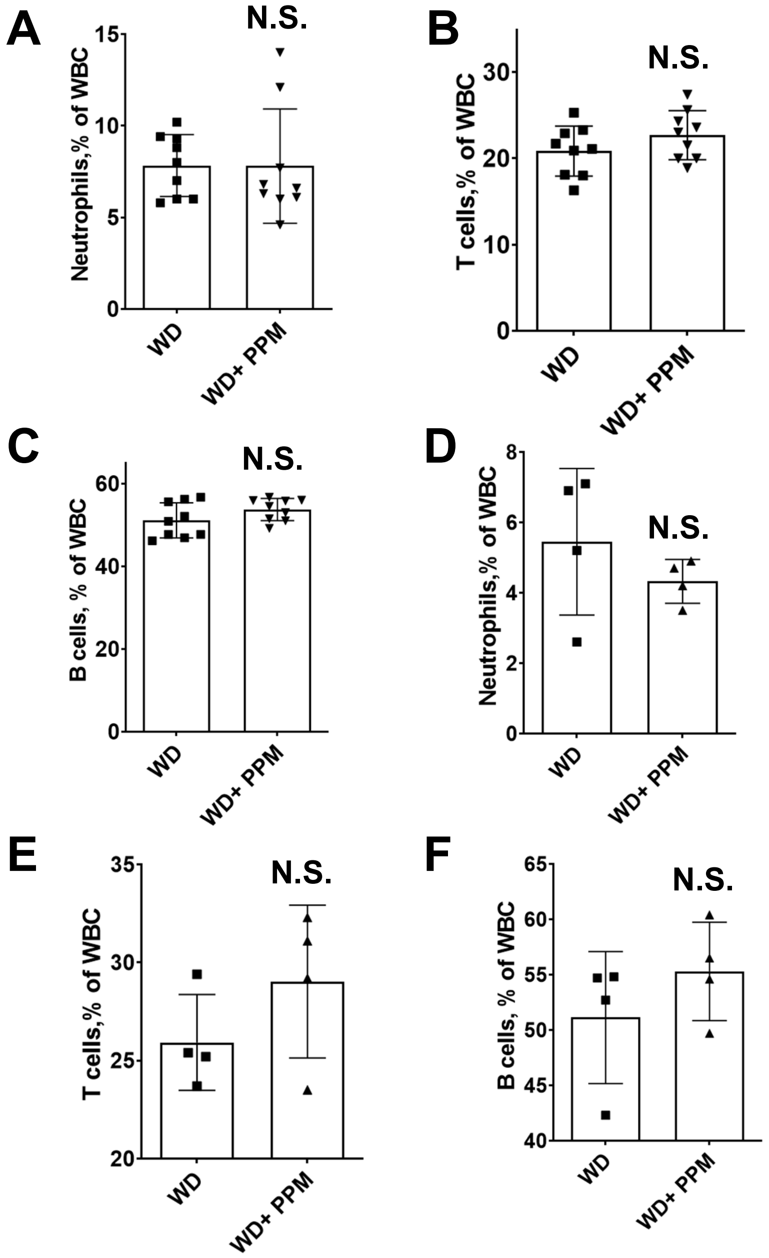


**Figure S11. PPM does not change the number of blood neutrophils, T cells, and B cells in female or male *Ldlr*$^{-/-}$ mice.** Female (A-C) or male (D-F) *Ldlr*$^{-/-}$ mice were treated with PPM or water alone and fed a western diet for 14 (A-C) or 16 (D-F) weeks. The number of blood neutrophils (A&D), T cells (B&E), and B cells (C&F) were then measured by flow cytometry as described in Methods. n=9 (A-C) or 8 (D-F) per group. Data are expressed as mean ± SEM, there is no significant difference between groups determined by using two-sided unpaired t-test.

### Figure S12

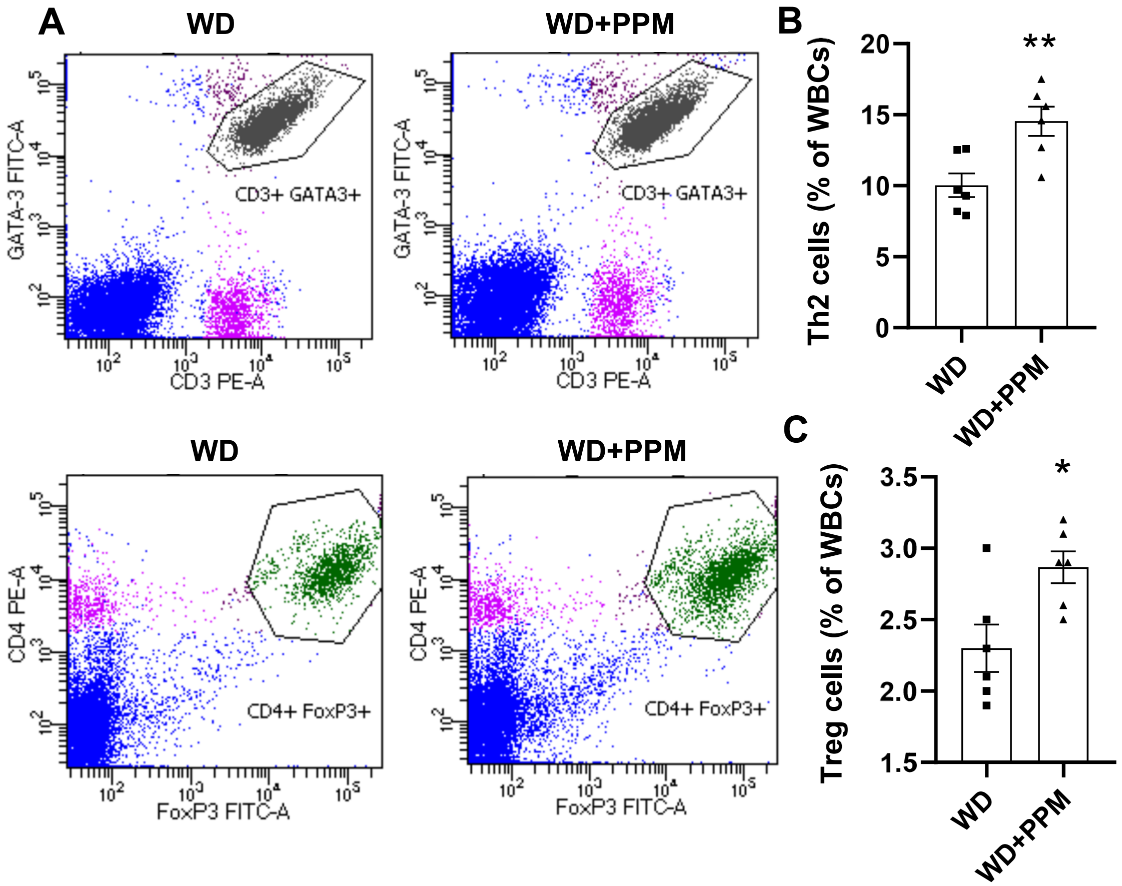


**Figure S12. PPM increases the number of blood Th2 and Treg cells in female *Ldlr*$^{-/-}$ mice.** A-B. The representative flow cytometry gating strategies for Th2 (A) and Treg (B) cells in peripheral blood of vehicle alone or PPM treated female *Ldlr*$^{-/-}$ mice fed a western diet for 14 weeks are shown. C. There was an increase in the number of blood Th2 cells (CD3$^+$CD4$^+$GATA-3$^+$CD8$^-$) in PPM versus water treated *Ldlr*$^{-/-}$ mice. D. PPM treatment increased the number of blood Treg cells (CD3$^+$CD4$^+$ FoxP3$^+$). C-D. n=6 in each group. Graph data are expressed as mean ± SEM; *P <0.05, ** P<0.01 by two-sided unpaired t-test.

#### Figure S13

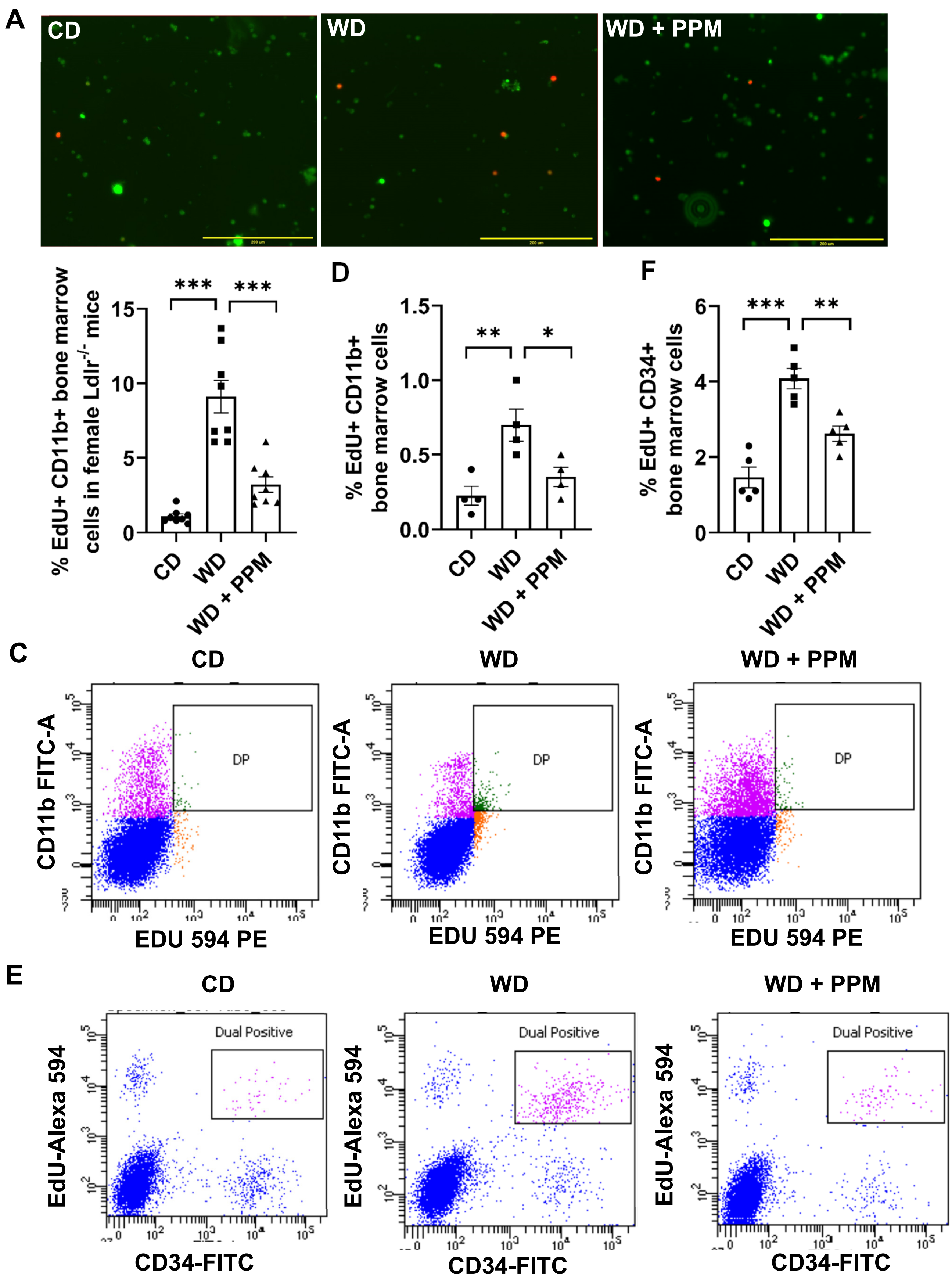
A
CD
WD
WD + PPM
B
% EdU+ CD11b+ bone marrow cells in female Ldlr-/- mice
***
***
CD
WD
WD + PPM
D
% EdU+ CD11b+ bone marrow cells
**
*
F
% EdU+ CD34+ bone marrow cells
***
**
C
CD11b FITC-A
DP
EDU 594 PE
E
EdU-Alexa 594
Dual Positive
CD34-FITC

**Figure S13. PPM decreases the proliferation of bone marrow monocytes and HSPC from female *Ldlr*$^{-/-}$ mice.** A-F. Bone marrow was isolated from female *Ldlr*$^{-/-}$ mice fed a chow or western diet for 16 weeks and treated with water alone or with PPM. A-D. Bone marrow cells were incubated for 24h with 5 μM EdU, and then stained with FITC-labeled CD11b antibody (green). EdU (red) was detected as described in the Methods. A-B. Dual EdU+ CD11b+ cells were detected by fluorescence microscopy (A) and quantitated (B). n=8 per group. Bar = 200 μM. Data are expressed as mean ± SEM; **** P<0.0001 by one-way ANOVA with Bonferroni's post hoc test. C-D. Dual EdU+ CD11b+ cells were measured by flow cytometry. n=4 per group. Data are expressed as mean ± SEM; *P <0.05, ** P<0.01 by one-way ANOVA with Bonferroni's post hoc test. E-F. Dual $EdU^{+}CD34^{+}$ cells were detected and quantitated by flow cytometry. n=4 each group, Data are expressed as mean ± SEM, *P <0.05, ** P<0.01 by one-way ANOVA with Bonferroni's post hoc test.
